# Anticancer immunotherapies transition postcapillary venules into high-endothelial venules that generate TCF1+ T lymphocyte niches through a feed-forward loop

**DOI:** 10.1101/2021.12.24.474088

**Authors:** Yichao Hua, Gerlanda Vella, Florian Rambow, Elizabeth Allen, Asier Antoranz Martinez, Marie Duhamel, Steffie Junius, Ann Smeets, David Nittner, Damya Laoui, Stefanie Dimmeler, Thomas Hehlgans, Adrian Liston, Guiseppe Floris, Diether Lambrechts, Pascal Merchiers, Francesca Maria Bosisio, Jean-Christophe Marine, Susan Schlenner, Gabriele Bergers

**Affiliations:** VIB Center for Cancer Biology, 3000 Leuven, Belgium; Laboratory of Tumor Microenvironment and Therapeutic Resistance, VIB Center for Cancer Biology, 3000 Leuven, Belgium; Department of Oncology, KU Leuven, 3000 Leuven, Belgium; Laboratory of Molecular Cancer Biology, VIB Center for Cancer Biology, Leuven, Belgium; Oncurious NV, 3000 Leuven, Belgium; Department of Imaging & Pathology, Laboratory of Translational Cell & Tissue Research and Department of Pathology, University Hospitals Leuven, KU Leuven, 3000 Leuven, Belgium; Department of Microbiology, Immunology, and Transplantation, KU Leuven, 3000 Leuven, Belgium; VIB Center for Brain and Disease Research, 3000 Leuven, Belgium; Department of Surgical Oncology, University Hospitals Leuven, KU Leuven, 3000 Leuven, Belgium; Lab of Cellular and Molecular Immunology, Vrije Universiteit Brussel, 1050 Brussels, Belgium; Myeloid Cell Immunology Lab, VIB Center for Inflammation Research, 1050 Brussels, Belgium; Laboratory for Translational Genetics, Department of Human Genetics, KU Leuven, 3000 Leuven, Belgium; Institute of Cardiovascular Regeneration, Goethe-University, 60590 Frankfurt am Main, Germany; Department of Immunology, University of Regensburg, 93053 Regensburg, Germany; Laboratory of Lymphocyte Signalling and Development, The Babraham Institute, Cambridge CB22 3AT, UK; Department of Human Genetics, KU Leuven, 3000 Leuven, Belgium

## Abstract

The lack of T-cell infiltrates is a major obstacle to effective immunotherapy in cancer. Conversely, the formation of tumor-associated tertiary-lymphoid-like structures (TA-TLS), which are the local site of humoral and cellular immune responses against cancers, are associated with good prognosis and have recently been detected in Immune Checkpoint Blockade (ICB)-responding patients. However, how these lymphoid aggregates develop remains poorly understood. By employing scRNA sequencing, endothelial fate mapping, and functional multiplex immune profiling, we demonstrate that antiangiogenic immune-modulating therapies evoke the transition of postcapillary venules into inflamed high endothelial venules (HEVs), which generate permissive TA-TLS-like lymphocyte niches with PD1^neg^ and PD1^+^TCF1^+^CD8 T cell progenitors that differentiate into GrzB^+^TCF1^neg^ TIM3^+^ PD1^+^ CD8 T effector cells. Tumor-HEVs require continuous CD8 and NK cell-derived lymphotoxin signals revealing that tumor-HEV maintenance is actively sculpted by the adaptive immune system through a feed-forward loop.

**In Brief:** Hua & Vella et al. reveal that effective antiangiogenic immunotherapy transitions postcapillary venules into inflamed high-endothelial venules (HEV), sustained by CD8 T and NK cell-derived signals through a feed-forward loop. Thereby, tumoral HEVs establish perivascular niches in which TCF1^+^ PD1+ lymphocytes expand and produce cytolytic PD1+ TIM3+ CD8 T cells that facilitate anti-tumoral immunity.

**Highlights:**

- High endothelial venule induction by anticancer immunotherapies generates perivascular immune niches permissive for TCF1^+^ PD1^+^ CD8 progenitor T cell expansion and production of TCF1^neg^ PD1^+^ TIM3^+^ CD8 effector T cells
- Tumoral high-endothelial venules exhibit characteristics of inflamed lymph node HEVs and postcapillary venules
- Postcapillary venules dynamically transdifferentiate into high-endothelial venules in tumors, which requires continuous signals from surrounding immune cells
- CD8 and NK cells drive tumoral high-endothelial venule formation during antiangiogenic immunotherapies in a feed-forward loop via lymphotoxin beta receptor signaling

## Introduction

Immunotherapy, specifically in the form of immune checkpoint blockade (ICB), has emerged as a major therapeutic modality in cancer and provided unprecedented benefits, although it is effective only in a minority of cancer patients (Hodi et al., 2010; Topalian et al., 2012). Unresponsive tumors to immunotherapies are commonly characterized by the absence of pre-existing Tumor Infiltrating Lymphocytes (TIL) and can be further subdivided into immune-excluded tumors, in which T cells have been attracted to the periphery of the tumor but fail to infiltrate, and immune-desert tumors, which are entirely devoid of T-cell infiltrates (Anandappa et al., 2020) In contrast, clinical responses to immunotherapy have been shown to correlate with pre-existing CD8 T cell infiltration, a T cell-instigated IFNγ-induced gene signature, and neoantigen burden (Ayers et al., 2017; Schumacher and Schreiber, 2015; Tumeh et al., 2014). Further, recent data provide evidence that checkpoint blockade and other immunotherapies may rely not on the reversal of exhausted T cells but the expansion and differentiation of intratumoral self-renewing TCF1^+^ progenitor CD8 T cells into cytolytic TCF1^neg^ TIM3^+^CD8^+^ T cells (Kurtulus et al., 2019; Siddiqui et al., 2019). Two different TCF1^+^ CD8 progenitor populations have been described, which can be distinguished by PD1 expression. In a progressing tumor, persistent exposure to cognate neoantigens can promote terminal T cell differentiation that restricts effector functions and induces negative immune regulators of T cell function such as the inhibitory receptor PD-1 (programmed cell death protein 1) (Wherry and Kurachi, 2015). Notwithstanding, PD1^+^TCF1^+^ CD8 T cells (pT_EX_) were able to mediate a proliferative response to immune checkpoint blockade and other immunotherapies by expanding at the tumor site and generating differentiated TCF1^neg^ TIM3^+^ PD1^+^ CD T cells (tT_EX_) that have cytolytic activity. Interestingly, TCF1^+^ progenitor T cells were found to reside in specific antigen-presenting-cell enriched niches within the tumor, while tumors that did not exhibit these niches displayed substantially fewer T cells (Jansen et al., 2019). Congruent with these observations, the presence of spontaneous tertiary lymphoid structures in tumors (TA-TLS) is commonly associated with good prognosis, augmented patient survival, and clinical responses to chemo-and immunotherapies (Gago da Graca et al., 2021; Sautes-Fridman et al., 2019). TLS display variably organized T- and B- cell aggregates, ranging from diffuse B- and T-lymphocyte clusters to segregated T cell-rich zones and juxtaposing B cell follicles with follicular dendritic cells (FDCs) and germinal center characteristics. Formation of these structures requires the disablement of the barrier and immunosuppressive properties of the angiogenic tumor vasculature that hinders the infiltration of lymphocytes, disables the cytotoxic function of PD1^+^-T cells, and triggers apoptosis of Fas-expressing CD8^+^T cells (Motz et al., 2014). Thus, TA-TLS commonly harbors specialized blood vessels reminiscent of high endothelial venules (HEV) that are uniquely poised to facilitate lymphocyte infiltration (Martinet et al., 2011). HEVs are normally found in secondary lymphoid organs (SLO) to transport naïve lymphocytes into lymph nodes where they get primed and educated. Due to their specific functions, HEV express high levels of lymphocyte adhesion molecules, i.e., the ligand of L-Selectin/CD62L, which is the homing receptor for T- and B-lymphocytes. L-Selectin ligands are sialomucins that entail sulfated mucin-like glycoproteins, including podocalyxin, endomucin, CD34, nepmucin, and GlyCAM-1 (rodent-specific) (Girard et al., 2012; Rosen, 2004). HEVs are very effective in retaining and transporting lymphocytes because their sialomucins undergo post-translational modifications by HEV-specific sulfotransferases and glycosyltransferases, including Carbohydrate Sulfotransferase 4 (CHST4)(Kawashima et al., 2005; Uchimura et al., 2005) and Alpha-(1,3)-Fucosyltransferase VII (FucT-VII) (Allen et al., 2017; Homeister et al., 2001; Maly et al., 1996).

Antiangiogenic and other vessel-normalizing therapies have been shown to reverse tumor-vascular abnormalities and reinstate T-cell transmigration by enhancing the expression of lymphocyte adhesion molecules (He et al., 2018; Jain, 2001; Rivera and Bergers, 2015; Schmittnaegel et al., 2017). This has provided a rationale for a combination of vascular-normalizing and immunotherapeutic approaches to sustain and improve therapeutic efficacy in cancer (Allen et al., 2017; Schmittnaegel et al., 2017). We have previously shown that antiangiogenic immunotherapy in the form of anti-VEGF/R plus anti-PD-L1 in pancreatic and breast cancers resulted in substantial T cell infiltration by inducing the generation of HEVs concomitant with the formation of T cell-enriched TL-like structures and improved tumor responses (Allen et al., 2017). Antiangiogenic immunotherapy induced lymphotoxin beta receptor (LTβR) signaling via the non-canonical NFκB pathway, which is required for HEV maintenance and expression of their characteristic L-selectin ligand PNAd in SLOs (Browning et al., 2005). In line, HEV formation was further enhanced by boosting lymphotoxin beta receptor (LTβR) in the tumor endothelium with an LTβR agonist or peptides of the LTβR ligand LIGHT which enhanced the influx of endogenous T cells and effective tumor killing in conjunction with immunotherapies (Allen et al., 2017; Johansson-Percival et al., 2017).

The induction of HEVs in tumors underscores the importance of the vasculature in regulating lymphocyte infiltration and may provide new avenues to induce and sustain efficacy to immunotherapies by overcoming the major restriction of T cell exclusion from the tumor microenvironment.

However, little is known about the generation and biology of HEVs in tumors. Here, we employed scRNA sequencing, fate mapping, and functional multiplex immune profiling to investigate the ontogeny, regulation, and function of tumor HEVs, the intimate relationship between TILs and HEV development and function, and provide evidence that HEVs form TA-TLSs encompassing lymphocyte niches permissive for TCF1^+^ CD8 cell expansion and differentiation in response to immunotherapies.

## Results

### Single-Cell RNA-Seq identifies common and disparate characteristics of TU-HEVs and LN-HEVs

To obtain insight into the nature and ontogeny of TU-HEVs during immune-modulating therapy, we first assessed the phenotypic commonalities and discords between HEVs in tumors and peripheral lymph nodes (LN-HEV) and tumor endothelial cells (TU-EC) by single-cell transcriptional profiling. Since the triple combination of anti-VEGFR2 (D), anti-PD-L1 (P), and lymphotoxin receptor beta agonist (Ag) had consistently induced multiple TU-HEVs with surrounding lymphocyte infiltrates in PyMT-bearing mice we reasoned to use this treatment strategy and model for analyzing TU-HEVs (Allen et al., 2017). We used two different approaches for single-cell RNA-sequencing (scRNA-seq) because HEVs are a rare subset of endothelial cells in tumors (approx. 0.2-4% of TU-EC). First, we isolated CD45^-^CD31^+^ MECA79^+^ TU-HEVs and peripheral LN-HEVs by cell sorting (FACS) using the standard MECA79 antibody that detects an HEV-specific epitope of the peripheral node addressin (PNAd) (Butcher et al., 2016), the mature multivalent L-selectin ligand for lymphocyte homing. Synthesis of this post-translationally modified PNAd, requires the HEV-specific fucosyltransferase Fut7, glycosyltransferase Gcnt1, and carbohydrate sulfotransferase Chst4 (Brulois et al., 2020). We also collected CD45^-^CD31^+^ MECA79^-^ tumor endothelial cells (TU-EC) from treatment-naïve PyMT tumors by FACS and subjected all three EC populations to full-length scRNA-seq by SmartSeq2 (**Figure 1A**). We obtained 490 high-quality cells (157 LN-HEV, 95 TU-HEV, 238 TU-EC) with a median library complexity of 3847 genes/cell. Unsupervised Louvain clustering and t-distributed stochastic neighbor embedding (t-SNE) identified a total of five EC subtypes reflecting homeostatic HEV (LN-HEV), inflamed HEV/PCV (TU-HEVs), blood EC (TU-ECs), mitotic EC, and very few lymphatic ECs. **(Figures S1A and S1B)**. Projection of LN-HEV, TU-HEV, and TU-EC populations into a two-dimensional tSNE space **(Figure 1B)** displayed phenotypic heterogeneity. We identified discriminative gene signatures for TU-EC, TU-HEV, and LN-HEV populations and further validated characteristic markers by immunohistochemistry on tumor and lymph node sections, respectively **(Figures 1C-1E**).

**Figure 1.**
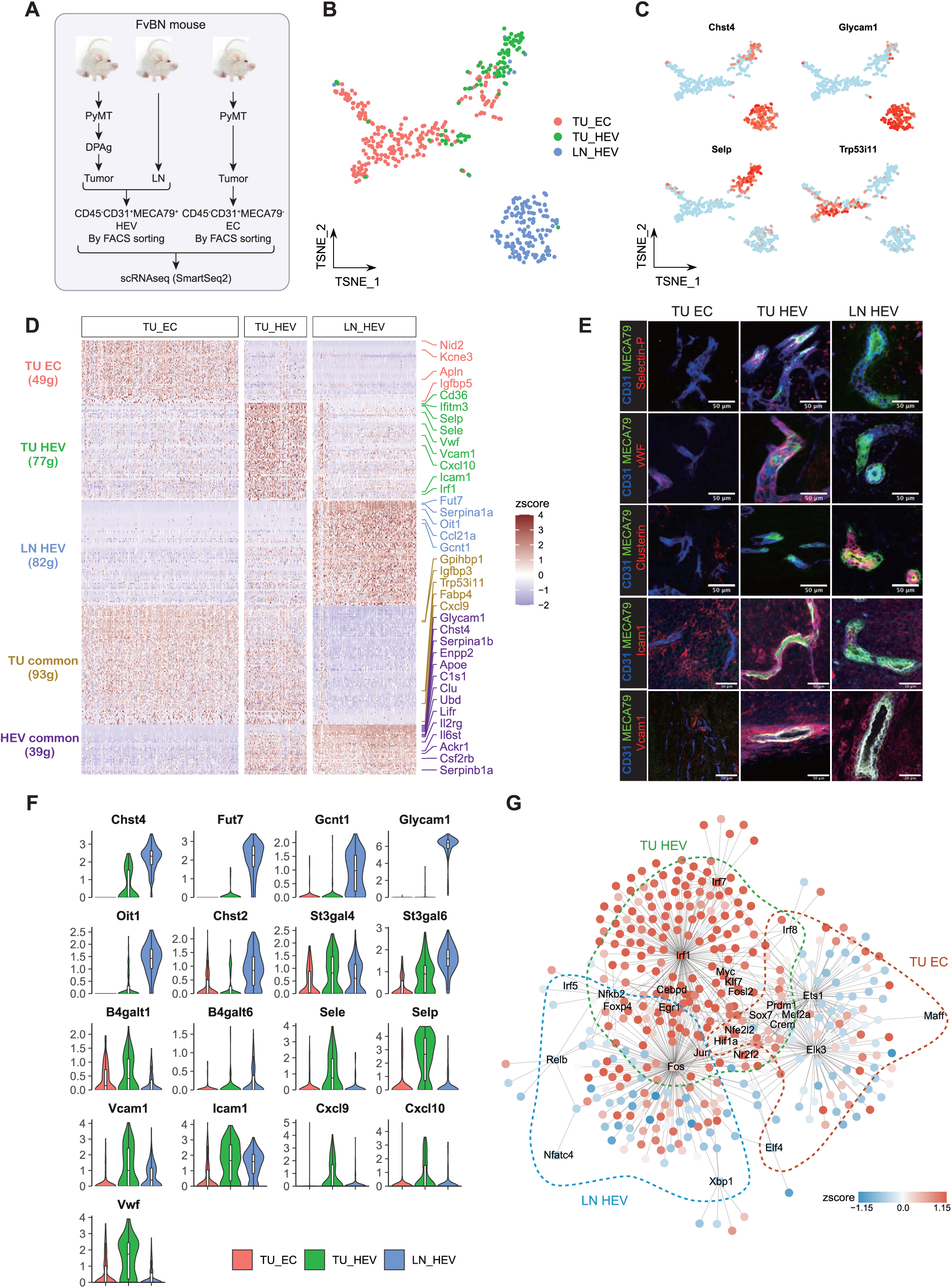
SmartSeq2 sequencing of TU-HEV, LN-HEV and TU-EC. (A) Study design, comparing transcriptomics of TU-HEV, LN-HEV and TU-EC. (B) tSNE plot of EC transcriptomes, colored by 3 different cell origins. (C) tSNE plots, colored by expression of representative marker genes (D) Expression levels of TU-EC, TU-HEV and LN-HEV specific and common genes (E) Immunofluorescence validation of HEV marker gene expression. Scale bars indicate 50 μm. (F) Violin plots displaying the expression of selected HEV and inflammation genes. (G) Gene regulatory network (GRN) predicted by SCENIC. The gene expression (round nodes) and regulon activity (square node) in TU-HEV is shown as node color.

Subsequently, differential gene expression between the different populations and associated Gene Set Enrichment Analysis (GSEA) **(Figures 1D and S1E)** confirmed that LN-HEVs displayed prominent transcriptional programs involved in glycoprotein synthesis and carbohydrate-based posttranslational modifications (Chst4, Chst2, Fut7, Gcnt1, St3gal6), as well as lymphocyte recruitment, adhesion, and diapedesis (e.g., GlyCAM1 (rodent-specific), Icam1, Vcam1, Ccl21a) (Brulois et al., 2020; Veerman et al., 2019) **(Figures 1D-F and S1C)**. As expected, a subset of TU-ECs proliferated (16%) and overexpressed endothelial tip cell markers (Kcne3, Apln, Nid2, and Trp53i110; **Figure S1B**), which is congruent with their overall angiogenic gene signature. TU-ECs, however, were devoid of the LN-HEV-specific gene expression profile **(Figures 1D-1F and S1C)**. TU-HEVs, like LN-HEVs, exhibited venule markers (Ackr1, Nr2f2) and expressed Glycam1, Chst4, Fut7, Clu, Oit1, and Gcnt1, but at lower levels, **(Figure 1F).** In addition, TU-HEVs displayed the highest levels of P-selectin (Selp), E-selectin (Sele), and Von Willebrand Factor (Vwf) compared to LN-HEV and TU-EC. Notably, all three factors are known to be increased during inflammation. Vwf is stored in the endothelial Weibel-Palade bodies and released during inflammation to promote platelet adhesion and hemostasis (Weibel, 2012). The cell adhesion molecules P-selectin and E-selectin bind the ligands PSGL-1 and ESL-1, respectively which are found on activated lymphocytes and myeloid cells. In concordance with these observations, TU-HEVs displayed elevated expression of a variety of additional genes implicated in interferon-regulated inflammation (e.g., Ifitm3, Irf1, Cxcl10) and antigen-processing compared to LN-HEV and TU-EC **(Figures 1D, 1F, 1G and S1E)**. Notwithstanding, TU-HEV and TU-EC also shared some transcriptional similarities (e.g., Gpihbp1, Igfbp3, Cxcl9) that were absent in LN-HEVs (**Figures 1D**). We then inferred the underlying gene regulatory code for our different EC populations using SCENIC (Aibar et al., 2017). As expected, the main transcription factors and associated gene regulatory networks (regulons) at play in each EC cell type showed commonalities but also unique differences. Besides Xbp1 and c-Fos, non-canonical NFκB, Relb, and NFκb2 transcription factors, implicated in lymphoid organogenesis, were activated in LN-HEVs, and to a lesser extent, in TU-HEVs, while the interferon-induced transcription factor Irf1 was highly activated in TU-HEVs confirming the GSEA results. As anticipated, transcription factors regulating angiogenesis (Maff, Ets1 and Elk3) were prevalent in TU-ECs **(Figures 1G and S1D)**. Taken together, these results demonstrate that TU-HEVs exhibit a hybrid phenotype of tumor endothelial cells and LN-HEVs with a prominent IFNγ gene expression signature.

### TU-HEVs exhibit features of inflamed postcapillary venules

Next, to validate our SmartSeq2 data, we assessed the presence of TU-HEVs in the context of the entire complexity of EC populations without pre-isolation by conducting transcriptional profiling of the entire tumor vasculature of naïve and DPAg- treated PyMT as well as E0771 breast cancers. We also interrogated the transcriptomes of endothelial cells from DPAg- treated PyMT tumors of mice exposed to DPAg plus anti-IFNγ treatment to assess whether IFNγ-induced inflammation instigated TU-HEV formation and phenotype. TU-ECs were isolated from the different tumors and conditions by FAC sorting (CD45^-^CD31^+^) and subjected to 10x Genomics which yielded a total of 9772 TU-ECs after quality filtering. We integrated the different data sets using the Harmony (Korsunsky et al., 2019) to overcome potential batch effects and projected the cells into a two-dimensional UMAP space **(Figures 2A and S2A).** For non-HEV EC subtype annotation, we used marker genes identified from published scRNAseq tumor endothelial datasets (Goveia et al., 2020). This resulted in ten endothelial clusters (M1-10) with varying abundance: arterial EC, two capillary subtypes (arterial and Aqp7), postcapillary venules (PCV), venous EC, lymphatics, tip cells, stalk cells, as well as ECs with a high interferon signature, and mitotic ECs **(Figures 2A and S2B).** We then aimed at identifying TU-HEVs with several “HEV core markers.” To do so, we identified Chst4 co-expressed genes (Pearson correlation) among highly-variable genes of the entire EC population **(Figure S2C)**. This resulted in the selection of Fut7, Glycam1, and Oit1, which besides Chst4, were also found to be highly expressed in LN-HEVs **(Figures 1D-1F and 2B)**. Oit1 is one of the top differentially expressed genes of HEVs compared to blood endothelial cells in LNs and Peyer’s patches (Lee et al., 2014). The activity of this core-HEV-signature (Fut7, Glycam1, Oit1, Chst4) was measured in each cell using the AUCell algorithm (Aibar et al., 2017). This resulted in 221 TU-HEVs (AUCell > 0.1) **(Figure 2C and S2D).** Among the different EC clusters, we found the majority of HEVs among postcapillary venules (PCV) with few amid the interferon and venule EC clusters and negligible numbers or none in the other EC subpopulations **(Figure 2D)**. By comparing the expression profile of TU-HEVs with other EC clusters, we identified three distinctive expression patterns of TU-HEVs comprising a PCV/venous, interferon/inflamed, and HEV-specific gene signature **(Figure 2E)**. These results confirm the previous SmartSeq2 analysis and reveal a phenotypic resemblance of TU-HEVs with LN-HEVs relating to their specialized production of lymphocyte adhesion receptors. Notwithstanding, TU-HEVs expressed elevated levels of Icam1, Vcam1, Sele, and Selp, which are induced during inflammation, enabling them to also recruit activated lymphocytes. Moreover, TU-HEVs, in part due to IFNγ-induced inflammation, exhibited prominent activation of the MHCI and MHCII antigen processing machinery that was devoid in LN-HEVs **(Figure S1E**). Taken together, these results confirm the SmartSeq2 data and provide further insight into the complexity of TU-HEVs.

**Figure 2.**
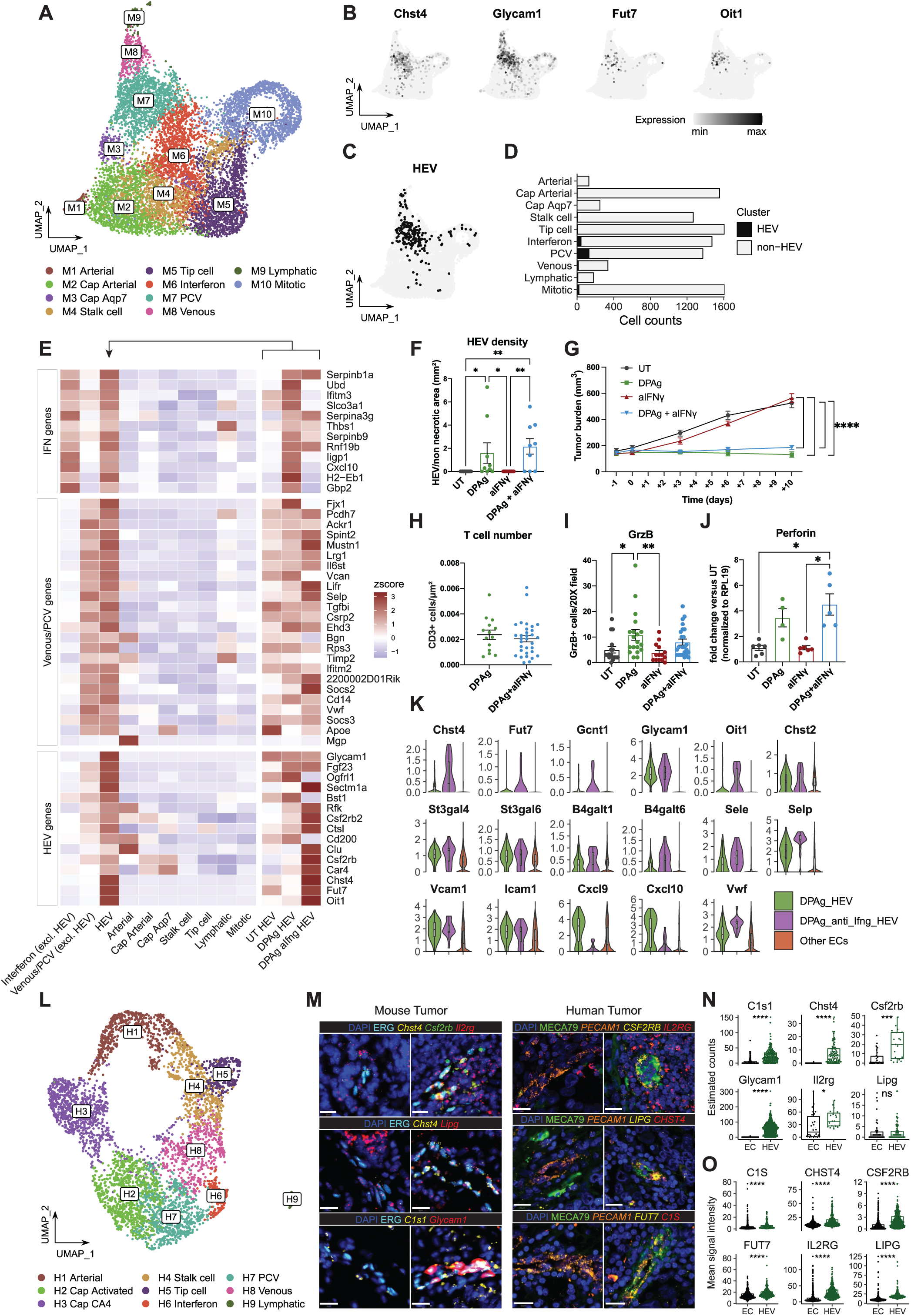
Characterization of the mouse and human tumor vasculature by droplet-based scRNAseq. (A) UMAP plot, colored by 10 EC subtypes identified by unsupervised clustering in mouse PyMT and E0771 tumors. (B and C) UMAP plots, colored by the expression of core HEV markers (B), or *in silico* selected HEV cells (C). (D) Fraction of HEVs per EC subtype. (E) Expression level of TU-HEV differentially expressed genes (DEGs) and core HEV markers, in tumor EC subtypes and TU-HEVs split by different treatments. (F) Quantification of HEV density of UT, DPAg, anti-IFNγ or the combination of DPAg and anti-IFNγ in PyMT tumors. HEV number was determined by immunofluorescence staining of CD31 and MECA79 on frozen tissues, normalized by the alive tumor area. N tumors: UT = 10; DPAg = 9; anti-IFNγ = 10; DPAg + anti-IFNγ = 9. The mean ± SEM are shown. Kruskal-Wallis test. (G) Tumor growth curves of treated PyMT-bearing mice. N tumors: UT = 11; DPAg = 12; anti-IFNγ = 12; DPAg + anti-IFNγ = 12. The mean ± SEM are shown. Two-way ANOVA. (H) Quantification of CD3^+^ T cells 50 μm^2^ around HEVs by immunofluorescence staining of DPAg or DPAg + anti-IFNγ-treated PyMT. N fields: DPAg = 14, DPAg + anti-IFNγ = 32. The mean ± SEM are shown. Mann-Whitney test. (I) Quantification of GrzB^+^ cells of PyMT tumors. N fields: UT = 10; DPAg = 9; anti-IFNγ = 10; DPAg + anti-IFNγ = 9. The mean ± SEM are shown. Krustal-Wallist test. (J) qPCR gene expression of perforin from total tumor lysate. RPL19 was used as housekeeping gene for gene expression normalization. N samples: UT = 6; DPAg = 4; anti-IFNγ = 6; DPAg + anti-IFNγ = 5. The mean ± SEM are shown. Krustal-Wallist test. (K) Violin plots visualize the expression of selected HEV and inflammation genes in HEVs in DPAg, DPAg + anti-IFNγ treated tumors and other ECs. (L) UMAP plot of ECs from human breast cancer samples, colored by 9 EC subtypes identified by unsupervised clustering. (M-O) Validation of selected conserved TU-HEV markers shared in both mouse and human datasets by RNAscope. In murine PyMT (N=1) and MC38 (N=1) tumors, ERG positive cells identify ECs and *Chst4*/ *Glycam1* positivity identifies HEVs (M, left panel), and the particle count of each RNA probe is quantified by QuPath (N). In human breast tumor (N=1), *PECAM1* positivity identifies ECs and MECA79 positive cells identify HEVs (M, right panel). Mean signal intensity of each probe in each *PECAM1*^+^ tile was measured (O) in QuPath. Wilcoxon test. Scale bars indicate 20 μm. Data are pooled from two independent experiments (F-J).

### IFNγ contributes to the disparity of TU-HEVs and LN-HEVs

The observed HEV-IFNγ responsive gene signature is likely to be the result of juxtapositioned activated lymphocytes known to secrete IFNγ, which raises the question about the implication of IFNγ in TU-HEV genesis. Yet, neither did DPAg immunotherapy alter the incidence of TU-HEV formation in PyMT-bearing mice **(Figure 2F)** when IFNγ was depleted, nor did it diminish the therapeutic effects of the DPAg treatment **(Figure 2G),** congruent with the observation that HEVs did not differ in their ability to endorse T-cell influx **(Figure 2H)** and that the T-cell cytotoxic proteins granzyme B and perforin were indistinguishably upregulated in both DPAg and DPAg + anti-IFNγ tumors compared to naïve tumors **(Figures 2I-J)**. Surprisingly, however, blocking IFNγ increased the expression levels of the HEV-specific core genes (e.g., Glycam1, Oit1, Fut7, Chst4), revealing that non-inflamed TU-HEVs become more reminiscent of homeostatic HEVs in lymph nodes (**Figures 2E, 2K)**. Indeed, the LN-HEV expression signature bore a significant resemblance to the one from TU-HEVs of DPAg plus anti-IFNγ treated tumors, while TU-HEVs of DPAg-treated tumors shared more similarities to that of an inflamed LN-HEV transcriptome (**Figures 2K and S2E) (Veerman et al., 2019).** In line with these results, IFNγ blockade increased expression of Glycam1, which binds L-Selectin^+^ naïve T-cells, and even further upregulated Selp and Sele mRNA levels. This is likely due to a negative feedback loop because IFNγ has been shown to reduce inflammation-induced expression of Selp and Sele in activated endothelial cells (Melrose et al., 1998). Thus, it is conceivable that although IFNγ does not change the ability of HEVs to facilitate lymphocyte infiltration, it may likely impact the leukocyte milieu in HEV-containing tertiary lymphoid-like structures.

### Murine and human breast cancer EC and HEV share a conserved gene expression signature

As several publicly available data sets of different human tumor types exist that encompassed endothelial cells, we set out to identify human TU-HEVs to test the phenotypic resemblance of their transcriptomic profile to our murine TU-HEVs. While we failed to detect any HEVs with our core Chst4 marker in these data sets, we obtained scRNA-seq data from 4621 breast cancer endothelial cells of a window-of-opportunity study (BioKey, NCT03197389) that had used 55 biopsies from pre-treatment and on-treatment early-diagnosed breast cancer patients obtaining one dose of anti-PD1 (pembrolizumab) (Bassez et al., 2021) **(Figure 2L).** We first clustered the human breast cancer endothelial cells into nine EC subtypes (arterial, two capillaries, PCV, venous, interferon, lymphatics, tip cell and stalk cells), which were similar to those of murine tumor endothelial cells entailing conserved markers for each mouse and human endothelial subcluster (Goveia et al., 2020): (**Figures 2L and S3C-E)**. Our attempt to identify HEVs *in silico* from the human datasets, however, only resulted in two Chst4^+^ endothelial cells found in the PCV cluster, which exhibited high Relb/NFκb2 activity levels like murine HEVs **(Figure S3A)**. We reasoned that the lack of HEVs in the human tumor samples was a combination of several incidents including the overall rarity of TU-HEVs, the small tissue acquisition from needle biopsies, and the mild single-cell isolation procedures accustomed for immune cells likely unable to free HEVs from their thick vascular shafts. Since the two Chst4^+^ endothelial cells were insufficient to derive conclusive information about human HEVs, we set out to identify a cell cluster that shared the expression profile and Relb/Nfκb2 activities with the two Chst4^+^ cells based on the diffusion components (Setty et al., 2019). This resulted in the identification of 90 human “HEV-like” cells, which shared over 20 conserved gene signatures with mouse TU-HEVs that were significantly increased in comparison to those of human PCVs and other non-HEV EC subtypes **(Figure S2F and S2G**). We selected the most conserved marker genes for both mouse and human TU-HEVs (e.g., Chst4, C1s1, Csf2rb, Il2rg) and confirmed by RNAscope that their expression in HEVs of murine tumor and human breast tumor tissues was significantly higher than in other endothelial cells **(Figures 2M-2O).** Taken together, these results reveal conserved transcriptional commonalities between murine and human tumor endothelial cells that are likely also shared among TU-HEVs.

### TU-HEVs are not terminally differentiated postcapillary venules

How do then HEVs arise in the tumor vasculature? To model potential TU-HEV differentiation trajectories in our 10x Genomics scRNAseq data set, we used Palantir (Setty et al., 2019) and scVelo (Bergen et al., 2020) (**Figure 3A**). We inferred the differentiation potential and gene expression diversity (developmental potential) of each cell using Palantir and CytoTRACE, respectively (**Figure S4A-B**) (Bergen et al., 2020; La Manno et al., 2018; Setty et al., 2019). Strikingly, we found a highly plastic “root cell” cluster in tumor endothelial cells from which most trajectories started and displayed distinct differentiation routes engendering five mature states defined as tip cells, capillary Aqp7, arterial, lymphatic and venous ECs **(Figure S4A)**. The latter seemed to arise by a continuum of developmental states with PCVs depicting an intermediate state. Chst4^+^ HEVs appeared within the PCV cluster, but like PCVs, were not recognized as a terminally differentiated cell type, indicating that PCVs and TU-HEVs bear some plasticity **(Figure S4B)**. In support, projection of naïve and DPAg-treated tumor endothelial cells revealed that the immune-modulating therapy significantly increased HEVs within the PCV cluster **(Figures 3A)**. CytoTRACE confirmed the observations of “root cells” having the highest DP (differentiation potential) while the four terminal states displayed a low DP, and PCVs and TU-HEVs exhibited an intermediate score **(Figure S4A).** These analyses infer that TU-PCVs have the necessary plasticity to convert to TU-HEVs and do so upon DPAg treatment.

**Figure 3.**
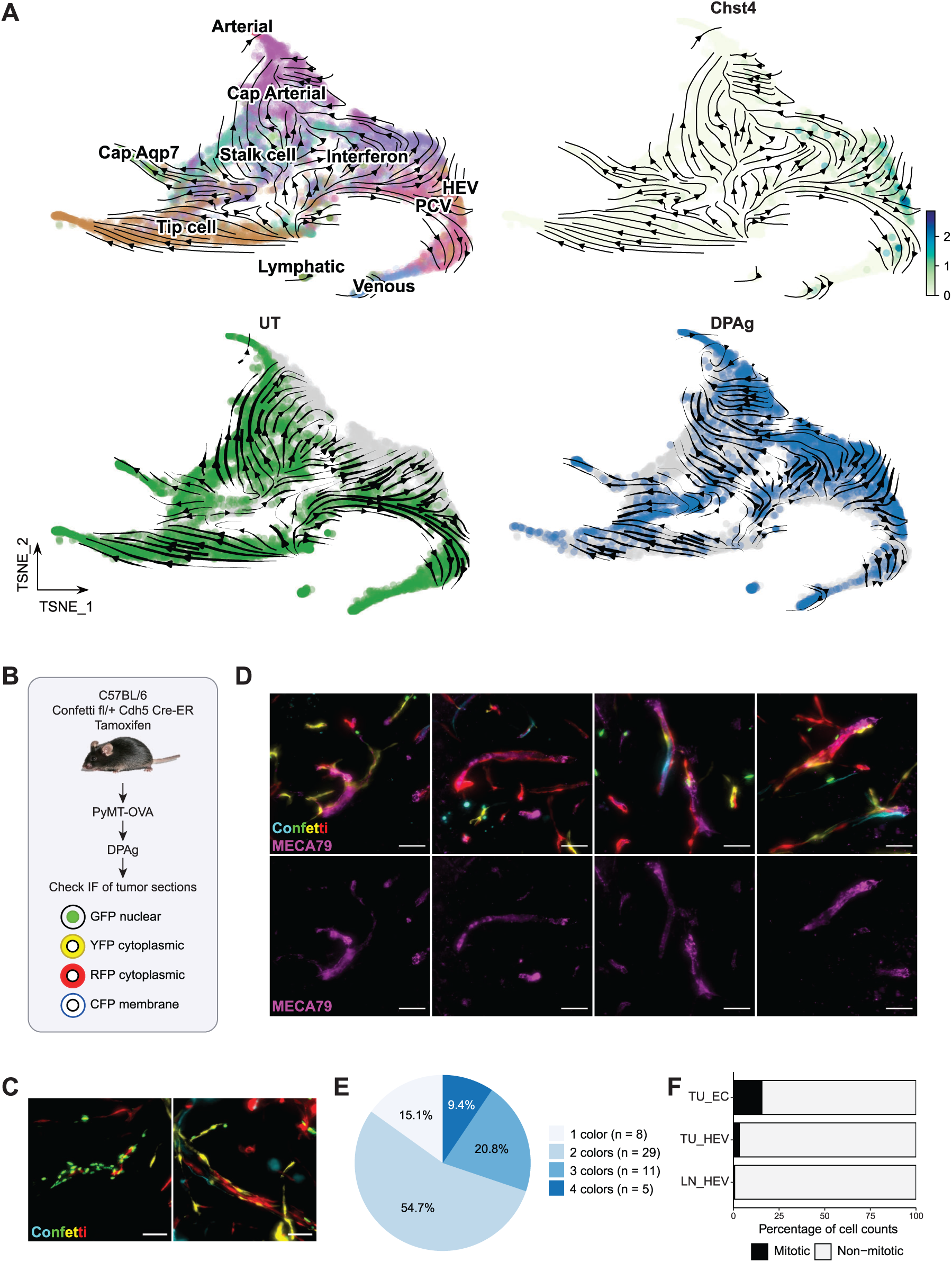
Ontogeny of TU-HEVs. (A) Differentiation trajectory of the entire tumor vasculature predicted by Velocyto/ScVelo, based on tSNE embeddings calculated by Palantir. Differentiation direction shown by arrows, in the entire datasets (top) and split by treatment groups (bottom). Dots color-coded for EC subtypes (top left), or Chst4 expression (top right) or treatment groups (bottom). (B) Study design of confetti tracing experiment. Recombination outcome after tamoxifen induction leads to expression of one of the following fluorescent proteins in endothelial cells: CFP (cyan fluorescent protein; plasma membrane), GFP (green fluorescent protein; nuclear), YFP (yellow fluorescent protein; cytoplasmic), and RFP (red fluorescent protein; cytoplasmic). (C) Representative images of ECs after tamoxifen induction. Blood vessels with single colored (left) or multiple colored ECs (right). Scale bars indicate 50 μm. (D) Representative images of MECA79^+^ HEVs of Confetti^fl/wt^Cdh5-CreERT2 PyMT-OVA tumor sections. Scale bars indicate 50 μm. (E) Pie chart showing the fraction of HEV vessels found with 1 to 4 colored ECs, respectively. (F) Fraction of mitotic cells in TU-HEV, LN-HEV and TU-EC from the SmartSeq2 datasets.

### TU-HEVs arise from postcapillary venules by metaplasia

Next, we asked whether TU-HEVs emanate from PCVs via metaplasia or by clonal expansion, of which the latter may suggest the presence of a progenitor or specific endothelial cell subtype. To investigate the lineage promiscuity of TU-HEVs at the level of individual endothelial cells, we took advantage of the R26R-Confetti tracer mouse model, which enables labeling and discrimination of individual cells with four nuclear-localized, membrane-targeted, or cytoplasmic fluorescent proteins in *cre* recombined cells (Manavski et al., 2018) (**Figure 3B**). Confetti^fl/wt^ mice were bred with mice expressing Cre-ERT2 (tamoxifen-inducible Cre-recombinase) under the regulation of the Cdh5 (cadherin 5 aka vascular endothelial cadherin) promoter (Confetti^fl/wt^ Cdh5-CreERT2) to induce endothelium-specific, tamoxifen-inducible conditional recombinase expression resulting in the exclusive, fluorescent labeling of ECs (Manavski et al., 2018). Upon injection of tamoxifen into Cdh5-CreERT2 Confetti mice, PyMT-OVA breast cancer cells were orthotopically injected, and mice were exposed to DPAg therapy to induce TU-HEVs (**Figure 23B and 3C**). We then assessed the color distribution of the four fluorescent markers in 53 MECA79^+^ TU-HEVs. Only 15% of HEVs comprised one color, while 85% of HEVs displayed two-four different colors indicating that the majority of HEVs did not arise from clonally expanding endothelial cells (**Figure 3D and 3E**). Notwithstanding, we observed areas of clonal expansion of TU-ECs (**Figure 3C**) displaying proliferating endothelial cells, which were, however, devoid of HEVs. These results indicate a metaplastic mechanism by which TU-EC transition into HEVs without the need for clonal expansion. In support of these findings, TU-HEVs depicted a very low proliferative rate of 3% in contrast to TU-ECs, of which 16% were mitotic (**Figure 3F**).

### TU-EC metaplasia into TU-HEVs is dependent on immunotherapy-induced signals

The rather prompt transformation of endothelial cells into TU-HEVs upon antiangiogenic immunotherapy raises the question as to which extent TU-HEVs depend on continuous therapy-induced signals. To evaluate the dynamics of HEV transformation, we treated PyMT and E0771 breast cancer-bearing mice with the triple immune-modulating DPAg therapy for 10-13 days and eight days, respectively, then stopped the treatment and followed tumor growth for another two weeks (**Figures 4A and 4C**). We evaluated the index of HEVs, and their functionality assessed by the morphology of MECA79^+^ cells and the number of surrounding lymphocyte infiltrates and conducted a detailed immune profile of tumors at the end of treatment and eight and 18 days after treatment cessation, as well as of naïve tumors as controls **(Figures 4B, 4D-4J, S4C-S4I)**. As we had previously shown, the immune-modulating therapy stabilized tumors and restricted tumor growth, which was associated with a substantial induction of TU-HEVs in both breast cancers after 10-13 days (PyMT) and 8 days (E0771) of treatment with 30-50% resembling mature, cuboidal-like MECA79^+^ endothelial structures (**Figure S4F**). The mature HEV phenotype also correlated with larger surrounding T cell infiltrates (**Figures S4F and S4H**). Zooming further into the lymphoid cell composition, TU-HEV induction was associated with an increase in intratumoral CD3, CD4, and NK cells as well as activated cytotoxic CD8 T cells as confirmed by IFNγ, TNFα and Granzyme B positivity, while the percentage of B-cells and CD4^+^FoxP3^+^ Tregs among CD4 cells were not substantially altered (**Figures 4E-4J**). Tumor-associated macrophages (TAM) did not follow this trend, being reduced in PyMT tumors and slightly increased in E0771 tumors during DPAg therapy **(Figures 4E, 4F and S4E)**.

**Figure 4.**
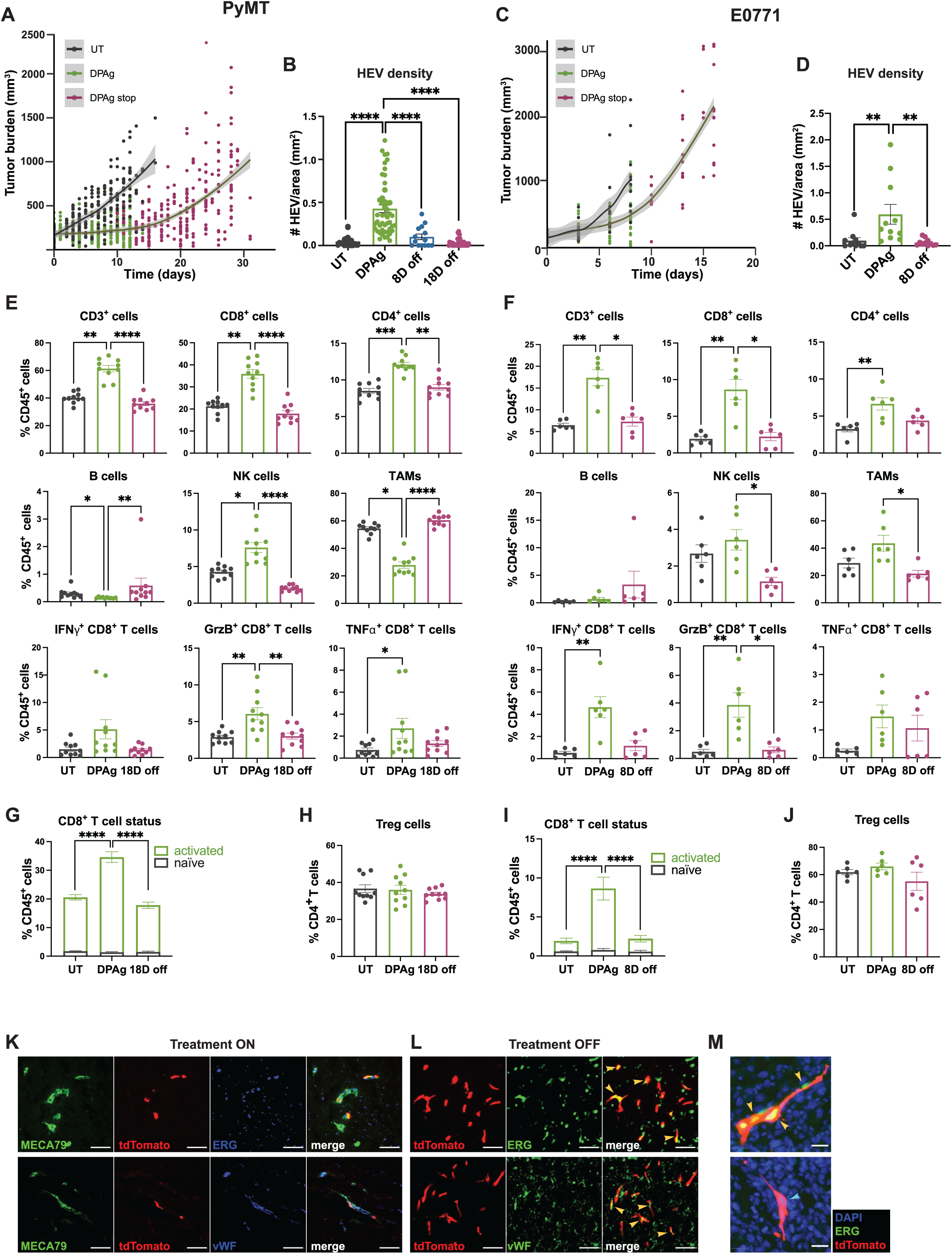
TU-HEVs dynamically arise upon immunotherapy and require continuous signals. (A and C) Growth curve of PyMT (A) and E0771 (C) tumors during and after DPAg treatment. N tumors: UT = 53; DPAg = 31; DPAg stop = 32 (A). N tumors: 11 for each cohort (C). 95% confidence interval (CI) of the curve is indicated by the grey line in A. (B and D) Quantification of TU-HEV density in PyMT (B) or E0771 (D) tumors by immunofluorescence tissue staining. HEV numbers are normalized by the total tumor area. N tumors: UT = 10; DPAg = 10; DPAg 8D off = 4; 18D off = 10 (B). N tumors: UT = 11; DPAg = 11; 8D off = 11 (D). The mean ± SEM are shown. Kruskal-Wallis test. (E-J) Immune cell characterization of PyMT (N=10 for each cohort) (E, G, H) or E0771 (N = 6 for each cohort) (F, I, J) tumors upon DPAg treatment or after treatment cessation (8D off or 18D off), by flow cytometry. CD62L staining was used to discriminate naïve (CD62L^+^) CD8^+^ T cells from the activated (CD62L^-^) CD8^+^ T cells (G, I). The mean ± SEM are shown. Kruskal-Wallis test. Only statistical differences UT vs DPAg, DPAg vs 18D off, and DPAg vs 8D off are shown. (K and L) Representative images of PyMT-OVA tumors from Chst4-tdT reporter mice upon DCAg treatment (treatment ON) (K) or after treatment cessation (treatment OFF) (L). Yellow arrow heads depict double-labeled cells. Scale bars indicate 50 μm. (M) Representative images of ERG^pos^ (yellow arrow) and ERG^neg^ (light blue arrow) tdTomato^+^ cells. Scale bars indicate 20 μm. Data were pooled from at least two independent experiments (A-J).

In contrast, when the immune-modulating therapy was terminated, tumors relapsed within a week, taking up the growth rate of tumors under naïve conditions **(Figure 4A and 4C).** In line with this observation, HEVs degenerated in tumors within one week and further declined two weeks after therapy termination with only very few and small MECA79^+^ HEV-like structures remaining in relapsed tumors **(Figures 4B, 4D, S4C, and S4D)**. The few TU-HEVs that persisted during treatment cessation, however, did not lose their ability to endorse lymphocyte infiltration into the tumor **(Figure S4I)**. Importantly, the decline of TU-HEVs was associated with a decrease in intratumoral CD4, NK cells, and activated CD8 T cells that reverted to levels similar to those observed in naïve tumors **(Figures 4E, 4F, 4G, 4I)**. Taken together, these data indicate that the immunomodulating therapy not only rapidly induced HEV formation and subsequent infiltration of activated lymphocytes but that it was also necessary to maintain the therapeutic conditions because HEVs and their TLS-like aggregates precipitously diminished upon the termination of immunotherapy, leading to tumor relapse.

### TU-HEVs dynamically arise upon immunotherapy and require continuous signals

Next, we inquired about the fate of TU-HEVs upon therapy cessation by genetically tracking TU-HEVs. We generated Chst4-CreER mice, which enables Cre-mediated gene deletion in HEVs, and bred them to Rosa26^LSL-tdTomato(tdT)^ mice (Chst4-tdT) in which Cre induces tdT expression in HEVs upon tamoxifen administration and remains expressed independently of cell fate. PyMT-OVA-bearing Chst4-tdT mice were treated with DCAg (a-CTLA-4 instead of a-PD-L1) for nine days and obtained tamoxifen starting six days after treatment initiation to induce tdTomato expression in TU-HEVs (on-treatment) **(Figure 4K)**. Then, the drug administration was stopped (off-treatment) **(Figure 4L)**. The on-treatment tumors were immediately analyzed, while the off-treatment group was analyzed 11 days after therapy termination when HEVs are diminished (off-treatment). Immunofluorescence staining of on-treatment tumors confirmed the appearance tdT-labeled MECA79^+^ TU-HEVs **(Figure 4K)**. About 73-84% of TU-HEVs expressed tdT likely due to their varying Chst4 expression levels and development of some TU-HEVs prior tdT labeling, while the majority of HEVs in lymph nodes were tdT-positive (**Figure S4J**). 11 days after treatment stop (off-treatment), tumors only contained a sparse population of tdT^+^ MECA79^+^ cells but displayed several tdT^+^/MECA79^neg^ cells at the tumor rim, where HEVs are commonly found, and some randomly distributed tdT^+^ cells within the tumor core. About 69% of tdT^+^ cells were endothelial cells at the tumor rim, as confirmed by Erg1 positivity. Further, IF staining revealed that some Erg1^+^ tdT-labeled cells expressed vWF, which is abundant in postcapillary venules **(Figure 4L)**. Surprisingly, we also observed some tdT^+^ cells in the tumor center, specifically after treatment cessation, of which only about 25% tdT^+^ cells appeared to be endothelial cells **(Figure 4L)**. These studies conclude that TU-HEVs require continuous signals to maintain their phenotype because they reverted to a non-HEV endothelial state as soon as the therapy-induced signals ceased. In addition, we found that HEVs not only transitioned back to a postcapillary venule/venous endothelium stage but that some endothelial cells even lost their endothelial phenotype when antiangiogenic immunotherapy ended (**Figure 4M**).

### CD8 T cells and NK cells induce immunotherapy-dependent HEV formation via the LT/LTβR axis

Since the presence of TU-HEVs strongly correlated with an increase in surrounding lymphocyte cell aggregates, we speculated that a feed-forward loop, instigated by paracrine signaling cues, is the underpinning mechanism of HEV-induced TLS formation. What are then the HEV-inducing signals, and from which immune cells do they emanate during the immunomodulating therapy? Since various immune cells are known to be involved in lymph node development and HEV maintenance in lymph nodes, and lymphocytes appear to regulate spontaneous HEV formation in tumors, together with the strong correlation of lymphoid cells surrounding therapeutically-induced TU-HEVs, we reasoned that recruited lymphoid cells might also provide the necessary HEV-inducing signals during immunotherapy (Colbeck et al., 2017; Johansson-Percival et al., 2017; Martinet et al., 2013; Moussion and Girard, 2011; Peske et al., 2015; van de Pavert and Mebius, 2010). In support, DPAg treatment of PyMT tumor-bearing Rag KO mice did not induce TU-HEVs and was rather ineffective in supporting the requirement of adaptive immune cells **(Figures S5I and S5J)**. Flow cytometric analysis of the PyMT and E0771-infiltrating immune cell repertoire revealed a negligible percentage of CD25^+^CD127^+^RORgt^+^ ILC3, the adult counterpart of Lymphoid Tissue Inducer cells (LTi) known to be involved in instigating lymph node development **(Figures S5D-S5H**) (van de Pavert and Mebius, 2010). Dendritic cell levels of pDC, cDC1, and cDC2 were also low and did not increase upon DPAg therapy **(Figures S5A-S5C)**. Thus, we first focused on the more prominent immune cell constituents in the tumor comprising CD8 T cells, CD4 T cells, NK cells, and macrophages (of which most were also CD11c^+^ ;data not shown). To interrogate the functional implication of these immune cells in TU-HEV genesis, we assessed TU-HEV generation in wildtype and CD4 T cell, CD8 T cell, NK cell, or macrophage-depleted PyMT tumors in response to DPAg therapy. While diminishing CD4 T cells or CSF1R^+^ macrophages did not affect HEV density in these tumors, depletion of CD8 T cells or NK cells reduced intratumoral HEVs and further diminished to levels of control IgG-treated tumors when both immune cell types were depleted **(Figure 5A)**. Tumors responded to the therapy independent of the presence of CD4 T cells or macrophages, displaying comparable HEV induction and elevated apoptosis to the wildtype counterparts **(Figures 5B and 5C)**. In contrast, CD8 or NK depletion reduced HEVs and the therapeutic anti-tumor effects, and combined CD8 T cell and NK depletion rendered tumors nearly ineffective to the immunotherapy which directly correlated with negligible HEV induction and a low apoptotic tumor index that was comparable to that of IgG control-treated tumors **(Figures 5B and 5C)**.

**Figure 5.**
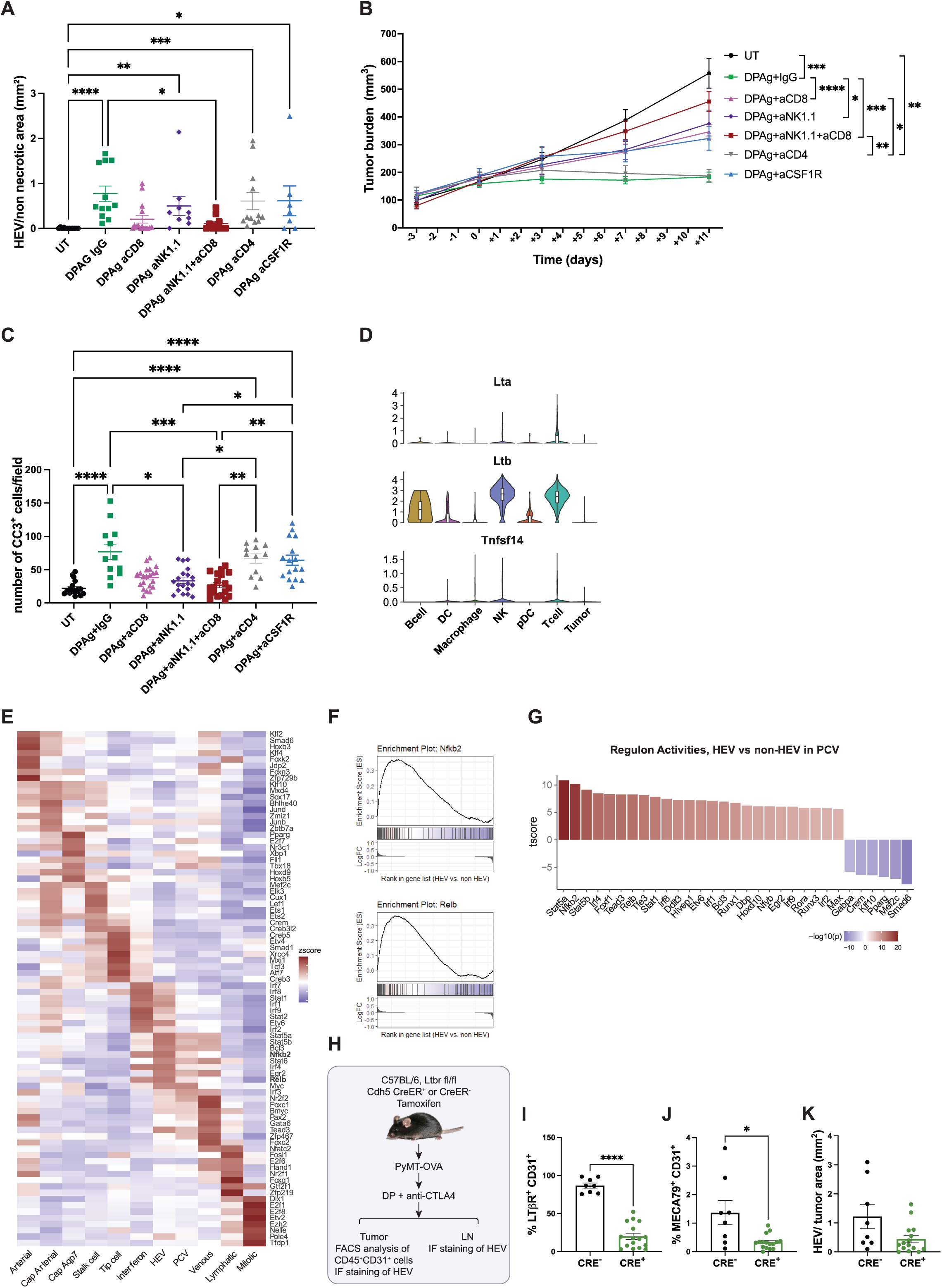
CD8 T-cells and NK cells induce immunotherapy-dependent HEV formation via the LT/LTβR axis. (A-C) TU-HEV density (A, N = 7-15), tumor growth curve (B, N = 6-15) and apoptotic index (C, N=12-21) of PyMT bearing mice treated under different conditions (treated with DPAg and depleted of CD8^+^ T cells, CD4^+^ T cells, NK cells or macrophages by administration of anti-CD8, anti-CD4, anti-NK1.1, anti-CSF1R, respectively). The mean ± SEM are shown. Krustal-Wallist test (A, C). Two-way ANOVA (B). (D) Violin plot of Lta (LTa), Ltb (LTb) and Tnfsf14 (LIGHT) expression in each immune cell cluster from the E0771 and PyMT dataset. (E) TF activities in each EC subtype predicted by SCENIC. (F) Gene Set Enrichment Analysis (GSEA) plots showing that Nfkb2 and Relb target genes were enriched in HEV cells compared to non-HEV cells. (G) Waterfall plot, comparing regulon activities of TU-HEVs and non-HEV ECs in PCVs predicted by SCENIC. (H) Experimental design of HEV evaluation in PyMT LTβR^ECKO^ mice. (Terai et al.) Quantification of intratumoral LTβR^+^CD31^+^ (I), MECA79^+^ CD31^+^ cells (J) by flow cytometry, and quantification of MECA79^+^CD31^+^ tumor vessels by immunofluorescence staining (K) of PyMT-OVA-bearing Cdh5-Cre^ER^×LTβR^fl/fl^ (CRE^+^) or control (CRE^-^) mice treated with DC101 + aPD-L1 + aCTLA-4 (DPC). N tumors: CRE^-^ = 8; CRE^+^ = 16 (I, J). N tumors: CRE^-^ = 8; CRE^+^ = 15 (K). The mean ± SEM are shown. Mann-Whitney test. Data are pooled from at least two independent experiments (A, C, D, I-K).

These results revealed that CD8 T-cells and NK cells are required cell constituents for the formation of TU-HEVs, which directly correlated with the beneficial effects of antiangiogenic immunotherapy. This raised the question as to whether these cells provide signals that are conducive for HEV conversion and maintenance. To identify the CD8 T cell and NK cell-derived factors that induce HEVs, we first conducted a NicheNet analysis (Browaeys et al., 2020) on our single-cell gene expression data to reveal the most prominent ligands that are highly expressed by CD8 T cells and NK cells and induce the expression of the respective DPAg-induced genes in ECs. IFNγ, lymphotoxin-α and β (LTα and LTβ) were the top-ranked ligands that were highly expressed by T cells and NK cells (**Figures 5D and S5L**). In line with this observation, using SCENIC, we found that the LT/LTβR -induced key transcription factors of the noncanonical NFκB pathway Nfκb2 and Relb exhibited increased activity congruent with enrichment of their target gene expression profile in HEV cells compared to non-HEV cells by Gene Set Enrichment Analysis (GSEA) of the scRNAseq data **(Figures 5E and 5F).** In addition, gene expression of the different participants that are involved in the LTβR – directed noncanonical NFκB pathway was also elevated in the HEV cluster compared to the non-HEV EC clusters **(Figure 5G).** These results confirm our previous data that antiangiogenic immunotherapy induced the LTβR pathway and that addition of an LTβR agonist further boosted HEV formation while an LTβR antagonist diminished it (Allen et al., 2017). To validate the scRNAseq and pharmacological data, we conditionally knocked out LTβR expression in endothelial cells by administering tamoxifen to Cdh5-Cre-ER-Ltbr^fl/fl^ mice. We then injected PyMT-OVA tumors and treated tumor-bearing mice with DC101, anti-PD-L1, and anti-CTLA-4 antibodies (DPC) to generate HEVs independent of the LTβR agonist (**Figure 5H**). Congruently, the treatment regimen significantly induced TU-HEVs wild-type PyMT-bearing mice but not in PyMT-bearing LTβR^ECKO^ mice as revealed by immunohistochemical and flow cytometric analysis **(Figure 5I-K).** Since the recombination efficiency among tamoxifen-treated Cdh5-CreER LTβR^fl/fl^ mice was, however, quite variable (50-90%), we tested whether the few TU-HEVs in some of the PyMT-Ltβr^ECKO^ mice arose from remaining LTβR wildtype or deficient endothelial cells. We bred the Cdh5-CreER-LTβR^fl/fl^ to the Rosa26^LSL-tdTomato(tdT)^ mice, which upon tamoxifen exposure, fluorescently labeled LTβR-knockout, but not wild-type, endothelial cells. These experiments confirmed that antiangiogenic immunotherapy induced MECA79^+^ TU-HEVs in LTβR-positive endothelial with few HEVs also arising from LTβR ko endothelial cells. **(Figure S5M).** Consistent with previous reports, we also observed that the LTβR knockout in endothelial cells substantially reduced HEV density in LN within two weeks (**Figure S5K**) (Veerman et al., 2019). All of the above results provide evidence that activated CD8 T cells and NK cells provide LTα_1_β_2_ lymphotoxins that induce TU-HEVs by activation of the LTβR and downstream noncanonical NFκB pathway in tumor endothelial cells. These signals are pivotal not only for the induction but also for the maintenance of TU-HEVs as well as LN-HEVs.

### TU-HEVs generate lymphocyte niches permissive for PD1^neg^ and PD1^+^ CD8 progenitor cells

TU-HEVs enable the infiltration of lymphocytes, but their composition and functionality is poorly understood. Here, we interrogated whether TU-HEVs can form lymphocyte niches that are permissive for T cell progenitor expansion and differentiation. Recent studies have shown that ICB therapy can promote differentiation of CD8 progenitor cells, rather than reversing dysfunction of terminally exhausted T cell (Kurtulus et al., 2019; Miller et al., 2019; Siddiqui et al., 2019). As such, progenitor cells appear to be the therapeutically relevant target of ICB therapy because they replenish and differentiate into Granzyme B (GrzB) (Miller et al., 2019) expressing CD8 T cell effector cells which also express the exhaustion marker TIM3 (Havcr2) (tT_EX_). Progenitor cells are commonly defined by expression of the transcription factor T cell factor 1 (TCF1) and the absence of TIM3 (**Figure 6C**) (Miller et al., 2019). Given that PD1^+^ TCF1^+^ TIM3^neg^ progenitor cells (pT_EX_) play a key role in CD8 T cell responses to ICB therapy, we investigated whether these cells also increase during antiangiogenic immunotherapy, accumulate in HEV^+^ lymphocyte niches and differentiate into PD1^+^TCF1^neg^TIM3^+^GrzB^+^ CD8 T cells (tT_EX_).

**Figure 6.**
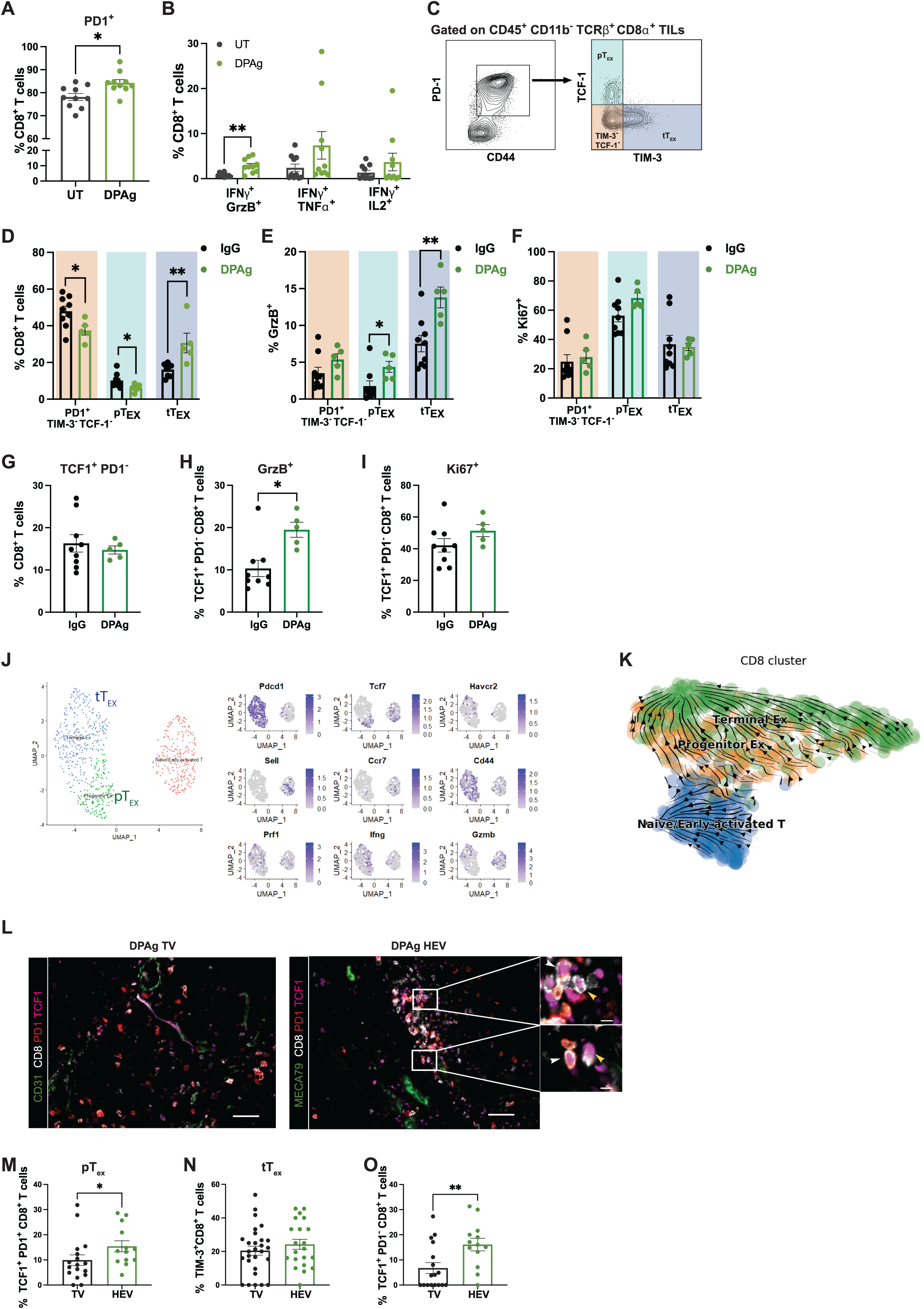
TU-HEVs generate lymphocyte niches permissive for PD1^neg^ and PD1^+^ CD8 progenitor cells. (A and B) Flow cytometry quantification of PD1^+^ CD8^+^ T cells (A) and CD8^+^ T cells co-expressing IFNγ-GrzB or IFNγ-TNFα or IFNγ-IL2 (B) in UT (N = 10) and DPAg (N = 10) treated PyMT tumors. The mean ± SEM are shown. (C) Gating strategy for intratumoral PD1^+^ TCF1^-^ TIM3^-^ cells, PD1^+^ TIM3^-^ TCF1^+^ pT_EX_ cells, and PD1^+^ TCF1^-^ TIM3^+^ tT_EX_ cells. (D and G) Flow cytometry quantification of PD1^+^ TCF1^-^ TIM3^-^ cells, pT_EX_ cells, tT_EX_ cells (D), and TCF1^+^PD1^-^ cells (G) in IgG (N = 9) and DPAg (N = 5) PyMT tumors. The mean ± SEM are shown. (E, F, H and I) Flow cytometry quantification of GrzB^+^ (E, H) and Ki67^+^ (F-I) PD1^+^ TCF1^-^ TIM3^-^, pT_EX_ and tT_EX_ cells(Sade-Feldman et al.) or TCF1^+^PD1^-^CD8^+^ T cells (H-I) from IgG (N = 9) and DPAg (N = 5) treated PyMT tumors. The mean ± SEM are shown. Mann-Whitney test. (J) UMAP plot showing CD8 T cell subsets in the PyMT dataset (left) and expression of selected genes (right). (K) Differentiation trajectory inference of CD8 T cell subtypes by Velocyto/ScVelo from the PyMT dataset. (L) Representative images of PyMT tumors from DPAg-treated mice. Scale bar indicates 50 μm. White head-arrows depict pTEX cells, yellow head-arrows depict TCF1^+^PD1^-^ cells. Scale bar in the selections indicates 10 μm. (M-O) Quantification of pT_EX_ (N = 13-17) (M), tT_EX_ (N = 21-27) (N) and TCF1^+^PD1^-^CD8^+^ T cells (N = 13-17) (O) 50 μm around HEVs or other TVs in DPAg-treated tumor sections. The mean ± SEM are shown. Mann-Whitney test (A-B; D-I; M-O). Data are pooled from at least two independent experiments (A-B; D-I; M-O). UMAP plots and trajectory derived from the PyMT dataset (UT and DPAg-treated tumors).

Flow cytometry analysis revealed that the majority of CD8 T cells expressed PD1 in PyMT and E0771 tumors and increased to about 80-90% during DPAg treatment (**Figure 6A and S6A**). Nevertheless, during treatment, a subset of CD8 T cells co-expressed elevated levels of IFNγ and tumor necrosis factor alpha (TNFα), or Granzyme B (GrzB), or IL-2, exhibiting a ‘‘polyfunctional’’ phenotype that has been associated with better tumor control (**Figure 6B, 6C and S6B**) (Miller et al., 2019; Spranger and Gajewski, 2015). Further flow cytometry analysis of PD1^+^ CD8 cells revealed 10% of pT_EX_ and 16% of tT_EX_ of which the latter nearly doubled during DPAg treatment. TCF1^neg^ TIM3^neg^ CD8 T cells were also present and comprised the majority of PD1^+^ CD8 T cells (**Figure 6D**). Only few TCF1^neg^ TIM3^neg^ CD8 T cells and pT_EX_ expressed GrzB, indicating that they are generally not cytotoxic but pT_EX_ were more proliferative than tT_EX_ and TCF1^neg^ TIM3^neg^ CD8 T cells and produced IFNγ (**Figures 6E,6F and 6J**). TCF1^+^ PD1^neg^ CD8 T cells comprised about 15-30% of CD8 T cells (**Figure 6A and S6A**). Although they did not appear to increase upon treatment, about double as many TCF1^+^ PD1^neg^ CD8 T cells were GrzB^+^ upon DPAg treatment (**Figures 6G, 6H**). These data indicate that PD1^+^ tT_EX_ and PD1^neg^ TCF1^+/lo^ cells constitute the vast majority of the GrzB-producing T cells.

To further investigate the phenotypic differences between the distinct CD8 cell populations of naïve and DPAg-treated PyMT tumors, we used scRNA-seq and compared the transcriptional profiles by uniform manifold approximation, projection (UMAP), and unsupervised clustering which gave rise to three major CD8 clusters that we also had identified by flow cytometry; i.e., naïve/early activated CD8 cells being PD1^neg^, CD44^neg^ Tcf7^+^/lo (encoding TCF1), Grzb^+^, CCR7^+^, and pT_EX_, and tT_EX_. The proximity of pT_EX_ and tT_EX_ on the UMAP confirmed the transcriptional similarity between these populations, consistent with the previously proposed lineage relationship between pT_EX_ and tT_EX_ (**Figure 6J**) (Siddiqui et al., 2019). Using Palantir + scVelo (velocyto) of the scRNA-seq analysis, we mapped a differentiation trajectory of these three CD8 populations. We found that the majority of pT_EX_ transitioned into tT_EX_ while only a very small subset of naïve/early activated T cells became effector CD8 T cells. As these analyses only gave inkling about the overall CD8 T cell composition and activity of the entire tumor, we next assessed whether these changes occurred in the TU-HEV-induced lymphocyte aggregates. Indeed, we found a higher degree of PD1^+^TCF1^+^ pT_EX_ and PD1^neg^TCF1^+^ CD8 T cells surrounding HEVs than tumor vessels of treated PyMT and E0771 tumors **(Figures 6L-6O, S6D, and S6E**). It is important to reiterate that tumor vessels enhanced T cell infiltration to a certain extent due to the vessel normalization effects driven by the antiangiogenic immunotherapeutic drugs **(Figure S4F**). We reasoned it more compelling to compare the immune cell-related effects of HEVs to “normalized” tumor vessels than to naive tumor vessels. Consistent with a previous report, tT_EX_ cells showed a broader distribution within the tumors and therefore did not display significantly different numbers between tumor vessels and HEVs of treated tumors (**Figure 6N and S6F**).

We then analyzed the CD8 T cell subsets in HEV^+^ MC38 tumors obtained from mice that had been treated with a combination of LTβRAg and anti-CTLA-4 antibodies. Here, we employed MILAN multiplex immunohistochemistry to identify the two TCF1^+^ CD8 progenitors and their differentiated TIM3^+^ CD8 progenies in relation to the presence or absence of TU-HEV, which required maintaining spatial resolution (Bolognesi et al., 2017). Digital reconstruction of the stained tumors using 6 cell markers revealed a confounding effect between the nature of the vessels and their location displaying multiple MECA79^+^ HEV clusters at the tumor rim and in peritumoral areas (**Figures 7A and S6G**). An expert pathologist (FMB) manually annotated the tumor bulk, tumor edge, and non-tumor areas. We then analyzed the numbers, proliferative and cytolytic index of the following CD8 T cell subpopulations: TCF1^+^ PD1^neg^ TIM3^neg^ CD8 T cells, TCF1^+^PD1^+^ TIM3^neg^ CD8 T cells (pT_EX_), TCF1^neg^PD1^+^ TIM3^+^ CD8 T cells (tT_EX_), and not-otherwise-specified (nos) CD8 T cells. We identified a total of 40,930 CD8 T cells in the four MC38 tumors of which 1.83% displayed pT_EX_, 10.84% TCF1^+^ PD1^neg^ T cells, 26.23% tT_EX_ cell while 61.09% comprised not-otherwise-specified (nos) CD8 T cells. We revealed that pT_EX_ and TCF1^+^ PD1^neg^ CD8 cells were more abundant in HEV-high areas while tT_EX_ cells spread further into the tumor and thus, prevailed more in HEV-low areas and tumor bulk (**Figure 7B and 7C**). Interestingly, pT_EX_ displayed a substantially higher proliferation rate in HEV-high areas than HEV-low locations, whereas TCF1^+^ PD1^neg^ CD8 cells and tT_EX_, independent of their location, were less proliferative (**Figure 7D**). In contrast, a high percentage of tT_EX_ expressed the cytolytic enzyme GrzB, particularly in the tumor bulk and in HEV-low areas but showed lower levels in HEV-high tumors (**Figure 7E and 7F**). Together, these results support the notion that HEVs support the formation of lymphocyte niches in which PD1^+^TCF1^+^ CD8 lymphocytes can self-renew and produce GzB^+^PD1^+^ CD8 T cell effector cells that migrate into the tumor and facilitate tumor killing in response to immunotherapies.

**Figure 7.**
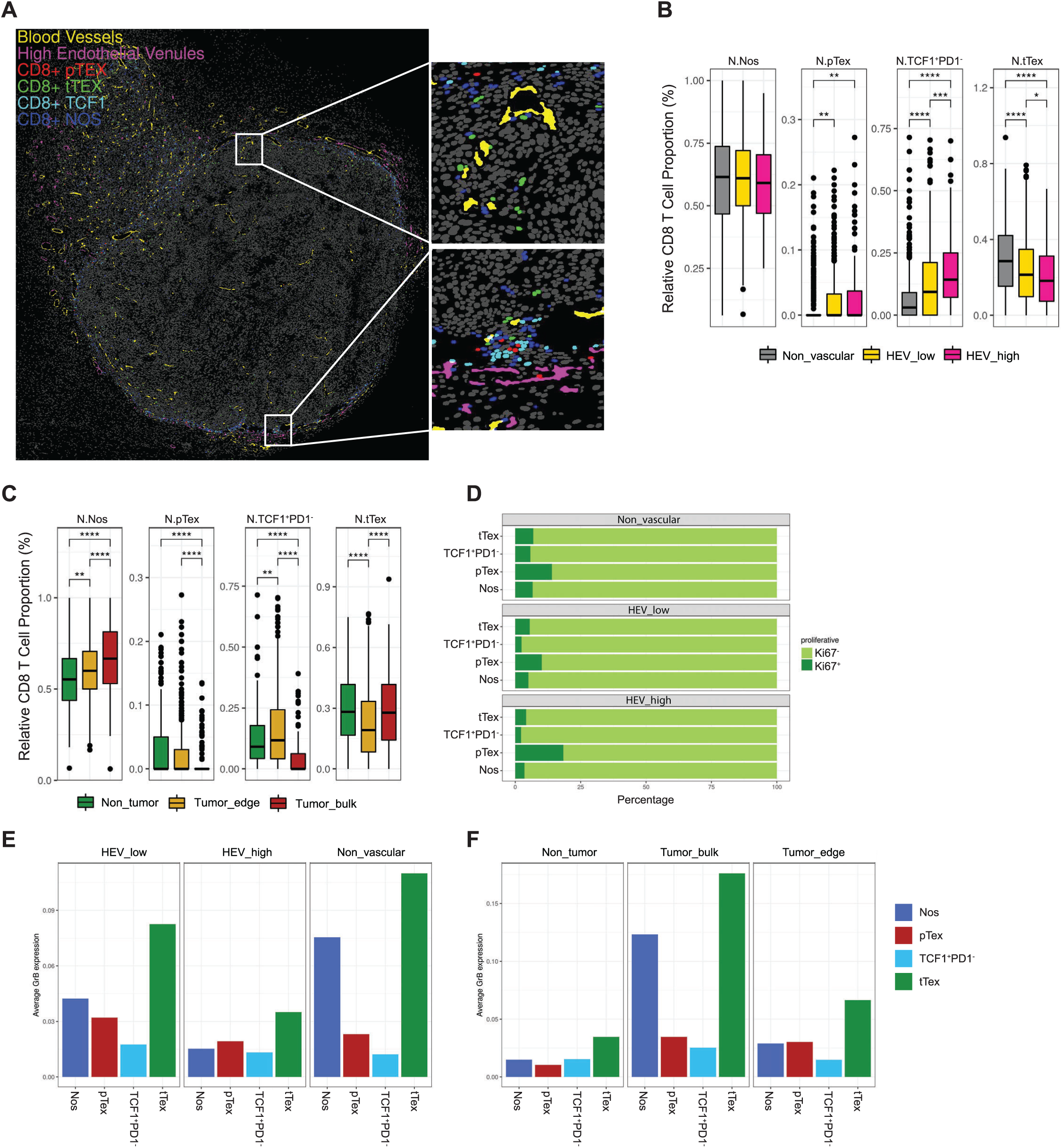
Immune and vascular landscape by MILAN. (A) Digital reconstruction of a representative MC38 tumor section stained with the MILAN multiplexing technique. Selected area depicts representative HEV-low area (upper square) and HEV-high area (lower square). CD8^+^ pT_EX_ (red) are TCF1^+^ PD1^+^ TIM3^-^; CD8^+^ tT_EX_ (green) are TCF1^-^ PD1^+^ TIM3^+^; CD8^+^ TCF1 (light blue) are TCF1^+^ PD1^-^ TIM3^-^. The remaining CD8 T cells are identified as CD8^+^ NOS (Not Otherwise Specified) (blue). (B and C) Boxplots indicating the relative proportion of different CD8^+^ T cell-types in the different tumor areas. (D) Fraction of proliferative (Ki67^+^) and non-proliferative (Ki67^-^) CD8^+^ T cells among the different subsets and in the different tumor areas. (E and F) Column charts depicting the average of GrzB expression of different CD8^+^ T cell-types in the indicated tumor areas. Data from B-F are derived from four MC38 tumors treated with anti-CTLA-4 and LTβR Agonist (both from Oncurious). Wilcoxon test (B and C)

## Discussion

Responses to anticancer immunotherapies are obstructed by the lack of intratumoral CD8 T lymphocytes. Thereby, the vasculature plays a fundamental role in controlling T cell infiltration. Our work demonstrates that induction of high endothelial venules in cancer does not just augment lymphocyte influx but is also critical for generating TCF1^+^ CD8^+^ T-cell niches in TA-TLS that, in response to anti-angiogenic ICB and ICB combination therapies, produce tumor-destroying GrzB-secreting TCF1^neg^ PD1^+^TIM3^+^ CD8 lymphocytes, which spread through the tumor. Notably, we found that TU-HEVs only form in an immunostimulating environment in which they require continuous signals emanating from activated lymphocytes and NK cells. In tumors, however, aberrant levels of proangiogenic factors promote vascular remodeling and growth, and endorse an immunosuppressive endothelial phenotype that limits T-cell influx and activity (Mazzone and Bergers, 2019; Motz et al., 2014). In contrast, inflammatory cues induce lymphocyte infiltration by elevating the lymphocyte adhesion molecules ICAM1 and VCAM1 on the endothelium. This parallels the vessel-normalizing effects of antiangiogenic therapy which abolishes the lymphocyte barrier and increases lymphocyte infiltration and improves anti-tumor immunity (Jain, 2001; Martin et al., 2019) Interestingly, in response to ICB, infiltrating, activated, INF-γ secreting lymphocytes have been shown to enhance vessel normalization (Tian et al., 2017). We have previously demonstrated that antiangiogenic ICB in the form of anti-VEGF/R plus anti-PDL1 not only increased vessel normalization and lymphocyte influx compared to single treatments, but also induced TU-HEV in breast cancer and pancreatic neuroendocrine (Lingscheid et al.) tumor-bearing mice (Allen et al., 2017). These data are congruent with our observation of a paracrine signaling loop between endothelial cells and activated lymphocytes, and further infer that vessel normalization may be an initial step in assembling additional perivascular activated lymphocytes, which upon a certain threshold, may yield endothelial conversion into high endothelial venules. HEV then amplify this feed-forward loop by further augmenting lymphocyte infiltration. A strong causal link between vessel normalization and HEV/TLS formation is further supported by a recent study in which PNET-bearing mice were treated with a tumor vessel-binding peptide of LIGHT (LIGHT-VTP), the ligand for LTβR and Herpes virus entry mediator (HVEM). Moderate LIGHT-VTP administration induced mature TA-TLS in direct correlation to the extent of vessel normalization but high doses of LIGHT-VTP impaired TA-TLS generation because it induced endothelial cell death (Johansson-Percival et al., 2017; Johansson-Percival et al., 2015). In summary, these data strengthen the proposition that vessel normalization and HEV induction are fundamental steps for TA-TLS formation.

With regard to TA-TLS composition, we predominantly observed T-cell aggregates with multiple MHCII+ positive myeloid cells around HEVs in tumors undergoing antiangiogenic immunotherapies. These resembled the dense aggregates of MHC-II^+^ APCs and CD8 T cells that had recently been discovered in human renal cell carcinomas (RCC) and were distinct from the typical B cell-enriched-identified TLSs found in RCCs, which did not exhibit closely interacting DCs and T cells (Jansen et al., 2019). Moreover, this study paralleled our results of increased TCF1^+^PD1^+^ progenitor CD8 T-cells (pTex) that give rise to terminally differentiated PD1+ TIM3+CD8 T cells (tTex) in these CD8 T-cell enriched areas (Jansen et al., 2019; Miller et al., 2019). Recently, both TCF1+ PD1^neg^ and TCF1+ PD1+ CD8 T cells have been described as self-renewing precursor populations that expanded and differentiated into Tcf1^-^ effector-like cells upon treatment with ICB immunotherapies producing a continuous pool of newly differentiated effector CD8^+^ T cells (Kurtulus et al., 2019; Siddiqui et al., 2019). We observed that pT_EX_, but not TCF1+ PD1^neg^ CD8 T cells, were more abundant and proliferative in HEV-high than HEV-low areas of treated PyMT, EO771 and MC38 tumors, whereas GrzB+ tT_EX_ were more enriched in the tumor bulk and in HEV-low tumor locations. The observation that tT_EX_ displayed augmented GrzB levels with increasing distance from HEVs is supportive of the suggestion that tTex surrounding HEVs are at an early stage of differentiation, just being produced by pT_EX_, and become cytolytically more active upoin maturation while they are migrating into the tumor bulk.

TA-TLS formation, similar to the development of secondary lymphoid organs (SLO), involves stepwise events and signaling cues between different immune and stromal cells (e.g., reticular fibroblasts, endothelial cells) (Johansson-Percival and Ganss, 2021 (Aoyama, 2021 #10997; Rodriguez et al., 2021; Sautes-Fridman et al., 2019). Lymphoid organogenesis is initiated by the crosstalk of lymphotoxin beta receptor (LTβR) expressing mesenchymal lymphoid tissue organizer (LTo) cells and hematopoietic lymphotoxin α1β2 (LTα1β2) expressing lymphoid tissue inducer (LTi) cells. LTo cells are believed to give rise to several mesenchymal cell constituents of SLOs like FRCs and the lymphatic and blood endothelium, including HEVs concomitant with expression VCAM1 and ICAM1 lymphocyte adhesion molecules and the LTi attracting chemokines Ccl19, Ccl21 and Cxcl13 (Johansson-Percival and Ganss, 2021).

Congruent with the notion that TU-TLS are believed to form via similar mechanisms, recent studies revealed cancer-associated fibroblasts (CAF) as surrogate LTo cells and CD8 T cells and LTα1β2+ B cells as LTi cells (Rodriguez et al., 2021). These aspects of SLO and TLS formation have been intensively studied, but the mechanistic underpinnings of HEV formation in SLO and TLS remains poorly understood (Ager, 2017). What has become apparent is that LTα1β2-producing CD11c+ dendritic cells (DC) are critical regulators of the HEV phenotype and function in lymph nodes because depletion of dendritic cells or genetic ablation of LTβR in endothelial cells severely diminished them (Moussion and Girard, 2011) (Girard et al., 2012). We observed few dendritic cells and many CD11c+ macrophages in HEV+ tumors but depletion of macrophages did not impair the ability of antiangiogenic immunotherapy to induce TU-HEVs. In contrast, we found that the presence of CD8 T cells and NK cells was required for HEV establishment during antiangiogenic immunotherapy as their depletion impaired HEV genesis in tumors. Similar results were observed with spontaneously arising HEVs in mouse tumor models (Peske et al., 2015). Importantly, the degree of HEV impairment directly correlated with the apoptotic index and anti-tumor effect, rendering antiangiogenic IBD therapy nearly completely ineffective in the absence of NK plus CD8 T cells. Using computational analyses, we uncovered the top signaling molecules emanating from lymphocytes that interact with tumor endothelial cells from treated tumors. Our studies revealed that lymphotoxins LTα/LTβ are the major drivers in immunotherapy-induced TU-HEV generation. In line with the requirement of NK and CD8 T cells in instigating TU-HEV, these cells exhibited the highest LT expression levels among other immune cell constituents, and endothelial-cell specific genetic deletion of LTβR severely reduced TU-HEV formation in ICB-treated tumors. Although LTβR signaling was the critical regulator for the maturation and acquisition of a fully mature HEV phenotype, we observed some immature HEVs derived from LTβR -deficient endothelial cells. These may have likely been generated by LTa3/TNFR signaling since this pathway is upregulated during inflammation and found to induce immature MECA79^+^ blood vessels in spontaneous HEV tumor models (Peske et al., 2015). Further, scRNA sequencing identified the non-canonical NFκB and downstream RelB/p52 pathway axis to be specifically and highly activated in TU-HEV. LTbR-induced activation of this pathway has been shown to be a critical regulator of lymphoid organogenesis and function (Jeucken et al., 2019) as LTβR deletion in endothelial cells abolished HEV formation subsequent LN homeostasis (Onder et al., 2013). IFN-γ inhibition did not reduce TU-HEVs but it promoted an HEV phenotype reminiscent of homeostatic HEVs in lymph nodes enhancing the levels of lymphocyte adhesion molecules for naïve lymphocytes (Veerman et al., 2019). Congruently, we found that the INF-γ gene signature in TU-HEVs resembled that of inflamed lymph node HEVs. TU-HEVs and inflamed HEVs upregulate P- and E-selectins to retain activated PSGL-1+ and ESGL-1+ T cells and myeloid cells indicative of the enhanced attraction of activated lymphocytes to the side. It is noteworthy that while lymphocytes, in general, express the genes of LTα and LTβ, and to a lesser extent LIGHT, only activated T cells have the lymphotoxin α_1_/β_2_ trimer bound to the cell surface, which underscores the link between an immunostimulatory environment and HEV formation (Browning et al., 2005; Browning et al., 1997).

How do tumor endothelial cells then develop into HEVs? TU-HEVs preferentially appear as hot spots close to the tumor rim, which is suggestive of specific vascular locations and endothelial cells from which HEVs can arise. This could further infer endothelial progenitors that give rise to HEVs. Intravital imaging of inflamed lymph nodes displayed clonal proliferation of HEV cells that acted as local progenitors to generate capillaries and HEV neovessels. It was suggested that lymph node homeostasis was likely achieved by the stochastic death of pre-existing and neosynthesized LN endothelial cells (Mondor et al., 2016). Recent scRNA sequencing and genetic lineage tracing identified a primed capillary resident regenerative population (CRP) that may function as progenitors to neogenesis of the blood vascular endothelium, including high endothelium in immune angiogenesis (Brulois et al., 2020). Using confetti multicolor fluorescent fate mapping and scRNASeq trajectory analysis of tumor endothelial cells and TU-HEVs, we found that TU-HEVs were not generated from a specific endothelial progenitor population by clonal expansion but rather by metaplasia from a common population of postcapillary venules. PCV conversion into HEVs occurred within days of antiangiogenic immunotherapy but also promptly reverted after therapy was stopped. These cells did not undergo apoptosis but reverted into postcapillary venules. In addition, a small subset appeared to further differentiate into non-endothelial mesenchymal cells, likely by EndMT, which reflects the plasticity of endothelial cells and response to antiangiogenic immunotherapies. The observation that lymphoid-derived signals promote PCV specialization to HEVs to accustom enhanced lymphocyte infiltration, and require a continuation of these signals to maintain in this specialized state, sets the stage for novel therapeutic opportunities that controllably induce HEVs and subsequent TCF1+ CD8 T cell containing TA-TLSs to improve current immunotherapies.

## STAR★METHODS

### KEY RESOURCES TABLE

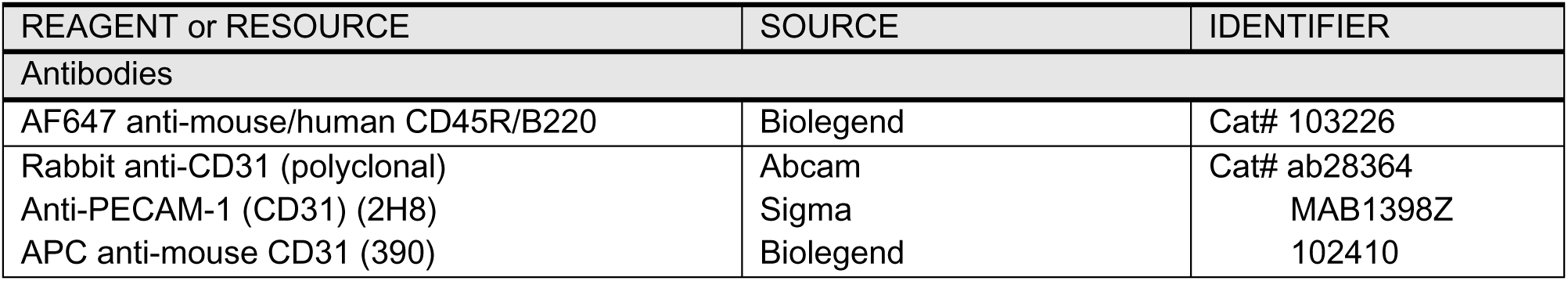
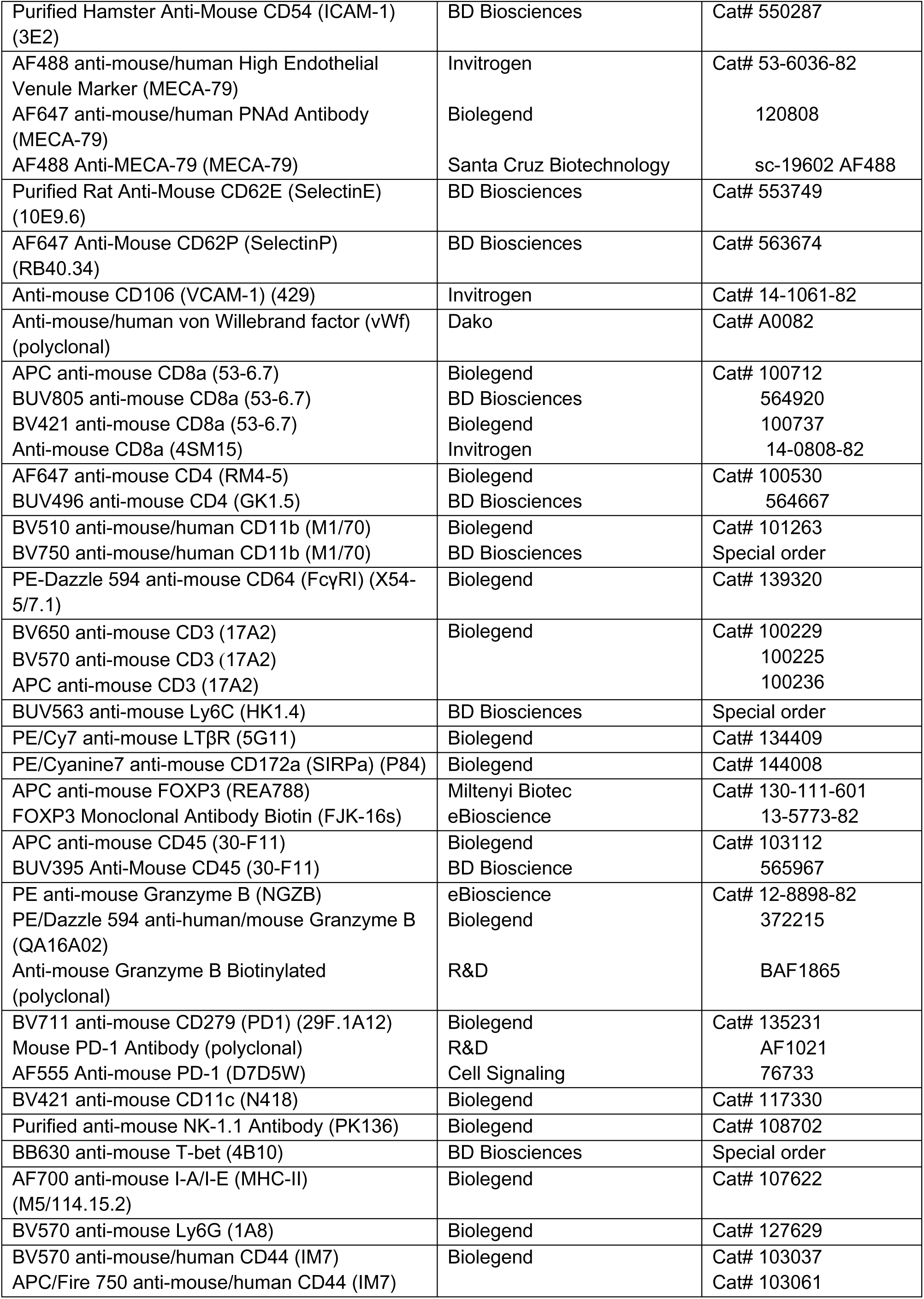
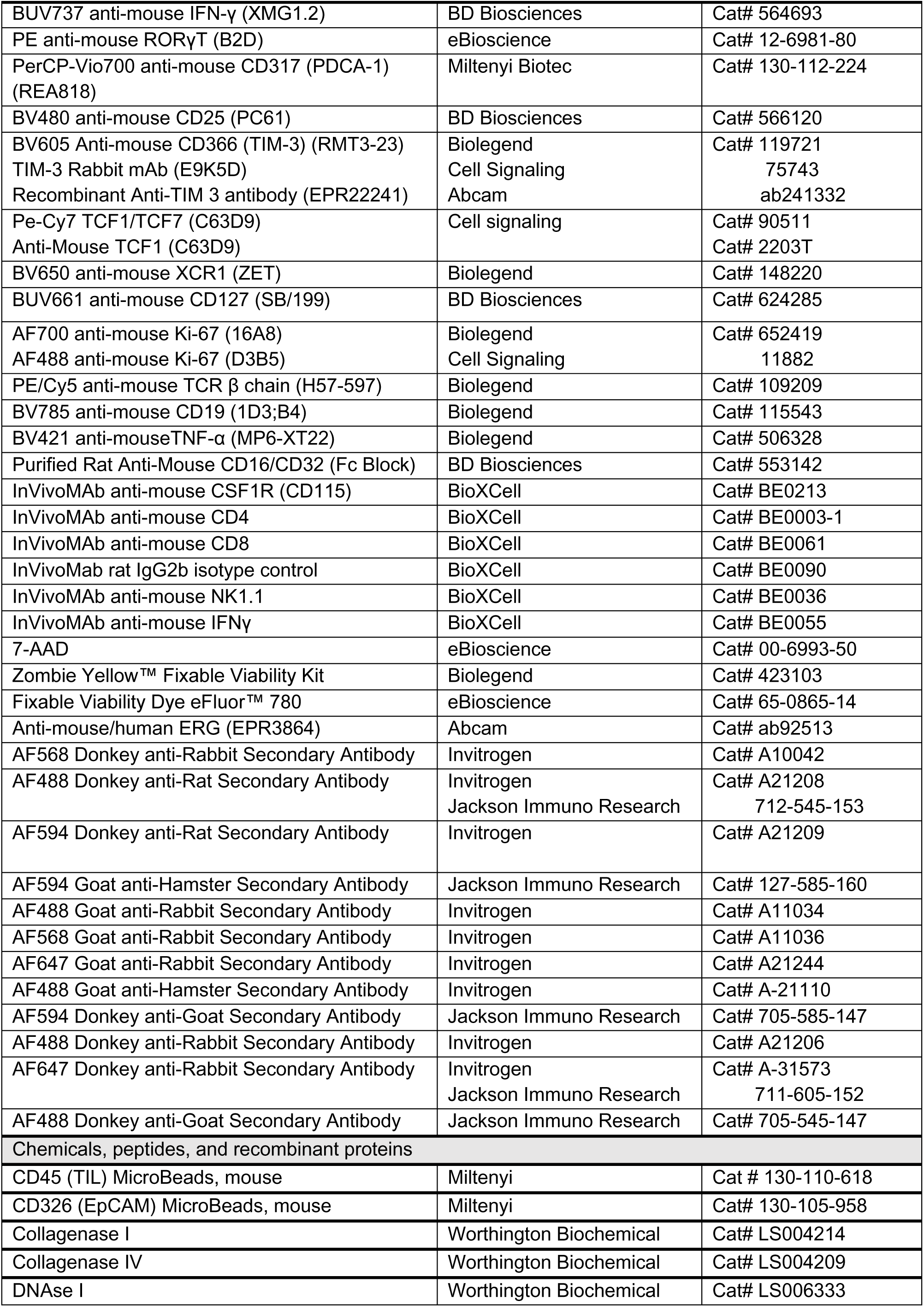
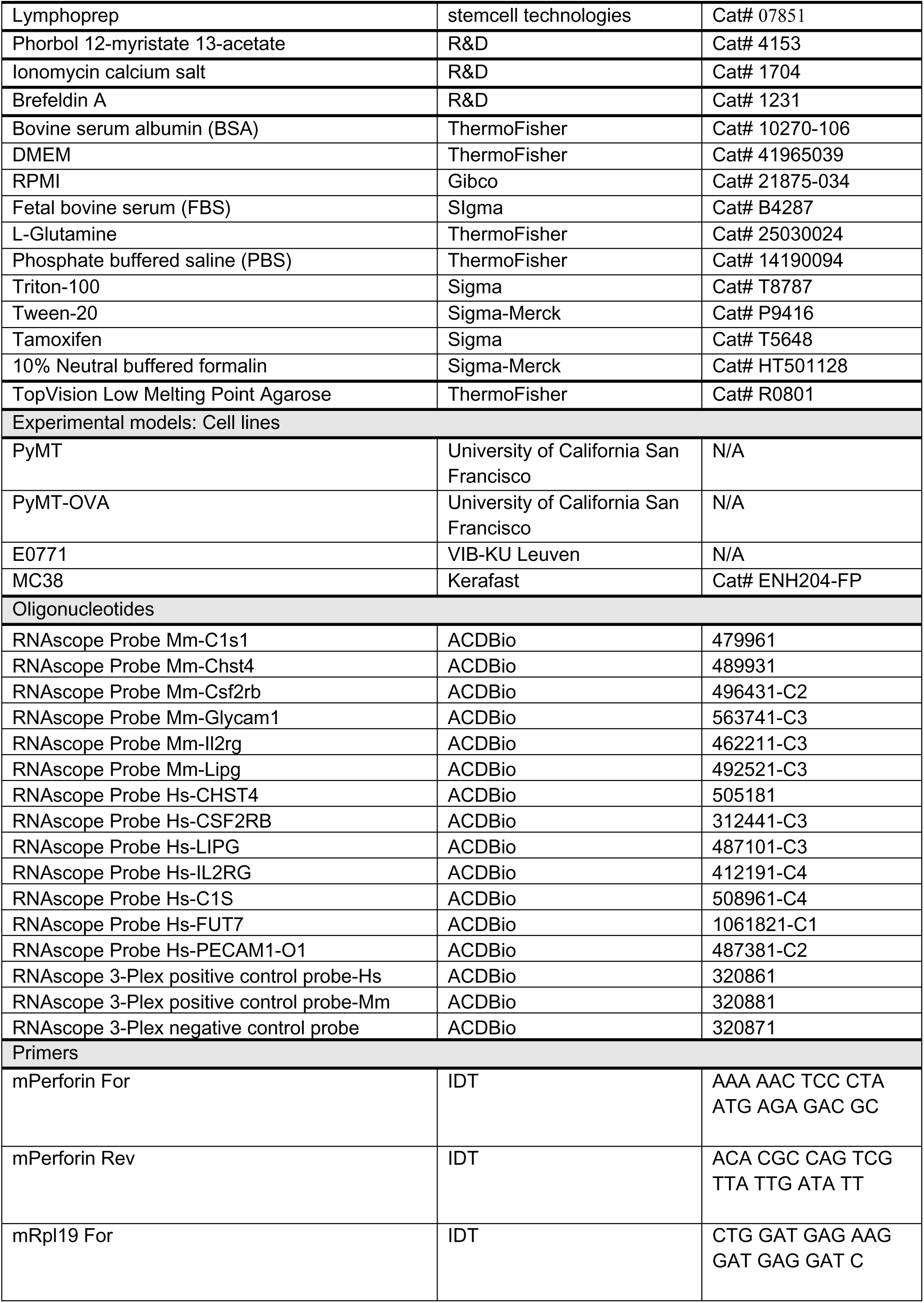
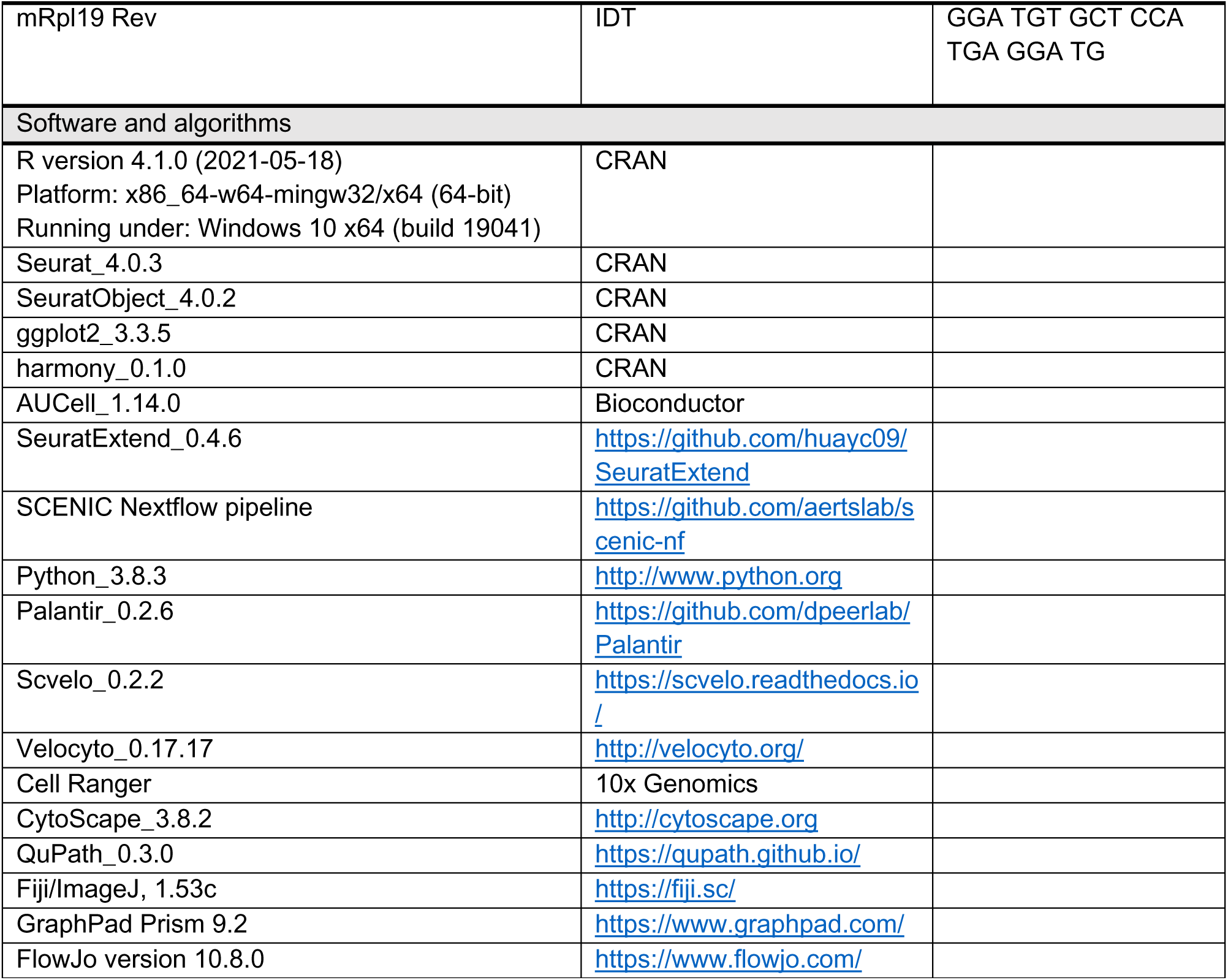

### LEAD CONTACT AND MATERIALS AVAILABILITY

Correspondence and requests for materials should be addressed to the. Lead Contact, Gabriele Bergers.

### EXPERIMENTAL MODEL AND SUBJECT DETAILS

#### Cell lines

The luminal breast cancer PyMT cell line was generated from FVB/N MMTV-PyMT mammary tumorwas akind gift from Dr. Zena Werb, University of California San Francisco). The PyMT-OVA cell line in the C57Bl/6 background was a kind gift of Dr. Max Krummel, University of California San Francisco.

The E0771 clone MC3B with metastatic potential to the lung (referred to as E0771) was kindly provided by Dr. Massimiliano Mazzone, VIB-KU Leuven, Belgium. The colon adenocarcinoma MC38 cell line was obtained from Kerafast. PyMT, PyMT-OVA, and MC38 cells were cultured with DMEM (Gibco) supplemented with 10% Fetal Bovine Serum (ThermoFisher), 1% glutamine, 100U/ml penicillin and 100U/ml streptomycin. E0771 were cultured with RPMI (Gibco) supplemented with 10% Fetal Bovine Serum (Gibco), 1% glutamine, 100 U/ml penicillin and 100 U/ml streptomycin. All cell lines were tested for mycoplasma every 3-5 passages.

#### Animal strains

Animal procedures were approved by the Institutional Animal Care and Research Advisory Committee of the KU Leuven (ECD 170/2016) and were performed following the institutional and national guidelines and regulations. C57BL/6, FVB/N mice were purchased from Charles River or obtained from the KU Leuven animal facility. Rag1 KO mice in an FVB/N background were in-house bred.

Conditional LTβR *knockout* mice (Cdh5-Cre^ER^×LTβR^lox/lox^ mice) were generated in a C57BL/6J background by intercrossing LTβR^lox/lox^ mice with the tamoxifen-inducible, endothelial cell-specific Cdh5-CreER mouse line. Cdh5-CreER^neg^×LTβR^lox/lox^ mice littermates were used as controls.

Confetti^fl/+^×Cdh5-Cre^ER^ mice in the C57BL/6J background (Manavski et al., 2018) were kindly provided by Dr. Stefanie Dimmeler from the Goethe University Frankfurt, Germany.

Chst4-Cre(ER)T2/iRFP mice were generated by ES-cell injection into fertilized blastocysts to generate chimeric mice, which were consequently bred with C57Bl6 albino animals to generate homologous recombinant mice. ES cells harbored a recombination of the following constructed transgene: The 5’ homology arm was comprised of 5836 bp of the Carbohydrate Sulfotransferase 4 (Chst4) promoter and 5’UTR, followed by the sequence encoding the first 6 amino acids (AA) of the CHST4 protein fused to AA 2-309 of iRFP670, a near infrared fluorescent protein that does not overlap with EGFP and tdTomato. An internal ribosomal entry site (IRES) preceded Cre-ERT, an SV40 poly-A signal, and a loxP site. An SV40 promoter drove expression of a neomycin resistance gene, followed by an SV40 poly-A signal, another loxP site, and a 3613 bp 3’ Chst4 homology arm with a third loxP site.

Chst4-tdT reporter mice (C57BL/6J) were generated by intercrossing the tamoxifen-inducible Chst4-Cre(ER)T2/iRFP mice with Rosa26-LSL-tdTomato reporter mice.

Conditional LTβR deletion (Cdh5-Cre^ER^×LTβR^lox/lox^ mice), the stochastic multicolor recombination in the endothelial cells (Confetti^fl/+^×Cdh5-Cre^ER^ mice), and the tdTomato labeling of TU-HEVs were achieved by intraperitoneal injections of Tamoxifen (Sigma-Aldrich) (2 mg/mouse/day) for 5 consecutive days prior to orthotopic implantation of PyMT-OVA cancer cells.

For all experiments, animals were maintained in individually ventilated cages in a room with controlled temperature and humidity under a 12-hour light/12-hour dark cycle with *ad libitum* access to water and food.

### METHOD DETAILS

#### Mouse trials

6 to 8-week-old FvB/N or C57Bl/6 female mice were anesthetized and a small incision in the skin of the lower abdomen was made to allow the orthotopic injection of 1×10^6^ PyMT or 2.5 x 10^5^ E0771 into each 4^th^ mammary fat pad. 1 x 10^6^ MMTV-PyMT-OVA were orthotopically injected in 9 to 14-week-old Chst4-tdT reporter mice, 8 to 11-week-old Cdh5-Cre^ER^ × LTβR^lox/lox^ mice, and 27-week-old Confetti^fl/+^×Cdh5-Cre^ER^ mice.

5 x 10^5^ MC38 cells were subcutaneously injected in the flank of 6 to 8-week-old C57Bl/6 female mice.

Treatment started when the tumors reached a volume of 100-150 mm^3^. Mice were treated with 20 or 40 mg/kg DC101(BioXcell), 10 mg/kg anti-PD-L1(BioXcell), 2 mg/kg LTβR agonist (Biorad), 2mg/kg LTβR agonist (Oncurious), 5 mg/kg anti-CTLA-4 (CD152) (BioXcell), 1 mg/kg anti-CTLA-4 (Oncurious), or 10 mg/kg IgG2b isotype control. All drugs were injected intraperitoneally (i.p.).

PyMT- and E0771-bearing mice received the DC101 + anti-PD-L1 + LTβR Agonist (DPAg) treatment twice a week for 10-13 days and 8 days, respectively.

PyMT-OVA-bearing Cdh5-Cre^ER^×LTβR^lox/lox^ mice received the DC101 + anti-PD-L1 + anti-CTLA-4 (BioXcell) (DPC) therapy every second day for 7 to 10 days.

PyMT-OVA-bearing Chst4-tdT reporter mice were treated with DC101 + anti-CTLA-4 (Oncurious) + LTβR Agonist (Biorad) every second day for 9 days.

PyMT-OVA-bearing Confetti^fl/+^×Cdh5-Cre^ER^ mice mice were treated with DPAg every second day for 9 days.

MC38-bearing mice were treated for three weeks, biweekly with 2 mg/kg LTβR Ag (Oncurious) and weekly with 1 mg/kg anti-CTLA-4 (Oncurious), starting 7 days after cancer cell implantation.

IFNγ was blocked by biweekly administration of 200mg/mouse of anti-mouse IFNγ (clone XMG1.2) (BioXCell) starting one day before the DPAg treatment until the end of the trial. For the immune cell depletion experiments, mice were injected i.p. with 0.5 mg anti-CD8 (clone 2.43), or anti-CD4 (clone GK1.5), or anti-NK1.1 (clone PK136), or anti-CSF1R (clone AFS98), or IgG2b (clone LTF-2) for 3 consecutive days before the start of DPAg treatment followed by weekly injection of 1mg of anti-CD8 or anti-CD4 or 0.5 mg of anti-NK1.1 or 0.5 mg of anti-CSF1R every second day until the end of the trial. All the depleting antibodies are purchased from BioXCell. Blood was collected before and after DPAg treatment to validate depletion efficiency by flow cytometry.

The t_0_ time point in the tumor growth curves indicates the treatment start. Unless differently specified, mice were sacrificed the day after the last treatment.

The tumor size was measured twice a week and the tumor volume was calculated using the following formula to approximate the volume of an ellipsoid: (width)^2^x length x 0.52.

#### Tissue Dissociation and Sample Preparation

For immune cell analysis, tumors were excised, mechanically minced and incubated with serum-free RPMI containing 3 U/ml Collagenase I (Worthington Biochemical), 133 U/ml Collagenase IV (Worthington Biochemical) and 10 U/ml DNAse I (Worthington Biochemical) and rotated at 37°C for 30 min. Enzymatic digestion was blocked by adding RPMI-1640 containing 2mM EDTA and 2% FBS. The remaining tumor aggregates were dissociated mechanically. The single cell suspension was then passed thought a 70µm strainer and red blood cells were lysed by incubation with 3 ml of ACK Lysis Buffer (15 mM NH_4_Cl, 10 mM KHCO_3_, 0.1 mM Na_2_EDTA) for 3 minutes at RT. For the immune characterization of PyMT and E0771 tumors upon DPAg treatment and at treatment cessation, and for the isolation of immune cells for the scRNAseq analysis, tumor-infiltrating lymphocytes were further enriched using Lymphoprep (Stemcell technologies) and density gradient centrifugation (800 x g for 30 min, without break or acceleration, RT). The interphase cells were collected and washed with PBS before flow cytometry staining.

For the analyses of TU-HEV, tumors were minced and incubated with enzymes A, D and R (Tumor Dissociation Kit, Miltenyi) in combination with mechanical dissociation by gentleMACS™ Dissociator (Miltenyi). Tumor endothelial cells were enriched by removing immune cells and tumor cells using the positive selection anti-mouse CD45 (TIL) MicroBeads (Miltenyi) and the positive selection anti-mouse CD326 (EpCAM) MicroBeads (Miltenyi). The negative fraction that contained the remaining endothelial cells was collected and washed with PBS before flow cytometry staining.

#### Flow cytometry

Single cell suspensions were blocked using 2.4G2 hybridoma supernatant or anti-CD16/32 blocking antibody (BD Biosciences). Samples were stained for 30 minutes on ice with antibody cocktails covering: CD31 (clone 390), MECA-79 (clone MECA-79), LTβR (clone 5G11), CD3 (clone 17A2), CD4 (clone GK1.5; clone RM4-5), CD8α (clone 53-6.7), CD11b (clone M1/70), CD11c (clone N418), CD19 (clone 1D3;B4), CD45 (clone 30-F11), CD62L (clone MEL-14), CD44 (IM7), CD64 (clone X54-5/7.1), CD172a (clone P84), Foxp3 (clone REA788), Granzyme B (clone NGZB; clone QA16A02), IFN-γ (clone XMG1.2), Ly6C (clone HK1.4), Ly6G (clone 1A8), MHC-II (clone M5/114.15.2), NK1.1 (clone PK136), PD-1 (clone 29F.1A12), PDCA-1 (clone REA818), TCR-β (clone H57-597), TNF-α (clone MP6-XT22), XCR1 (clone ZET), TCF-1 (clone C63D9), Ki67 (clone 16A8), NKp46 (clone 29A1.4), T-bet (clone 4B10), CD25 (clone PC61), CD127 (clone SB/199), and TIM-3 (clone RMT3-23) from Cell Signaling, Biolegend, BD Bioscience, eBioscience and Miltenyi Biotec. Dead cells were excluded using the Fixable Viability Dyes (eBioscience, Biolegend) or 7AAD (eBioscience).

For intracellular cytokine staining, lymphocytes were plated in U-bottom 96-well tissue-culture plates in complete RPMI containing phorbol 12-myristate 13-acetate (500 ng/ml Bio-Techne), ionomycin (750 ng/mL, Bio-Techne), and brefeldin A (2 μg/ml; Bio-Techne) for 4 hours at 37°C. Cells were blocked with 2.4G2 hydridoma supernatant in combination with mouse IgG (ThermoFisher) for 20 min at 4°C. Cells were fixed with 2% formalin and permeabilized by Foxp3/Transcription Factor Staining Buffer Set (ThermoFisher) before intracellular staining. For transcriptional factor staining, single-cell suspension was fixed and permeabilized by Foxp3/Transcription Factor Staining Buffer Set (ThermoFisher) before intracellular staining. TIL data were acquired on a BD FACSymphony or BD FACSCanto II (BD Biosciences). Conventional CD4 T cells and Treg cells were sorted by a BD FACS Aria Fusion (BD Biosciences).

Endothelial cell-related data were acquired on a BD FACSCanto II, CD31^+^ MECA79^-^ ECs and CD31^+^MECA79^+^ HEV cells were sorted with a BD FACSAria III (BD Biosciences). Results were analyzed with FlowJo version 10.8.0 (Becton Dickinson, Ashland).

#### Gene expression analysis

RNA was extracted from tumor lysates using the TRIzol^TM^ reagent (Life Technologies) and the PureLink™ RNA Mini Kit (Thermo Fisher Scientific) according to the manufacturer’s instructions. Pre-amplified cDNA from tumor lysates was made using the SuperScript^TM^ III First Strand cDNA Synthesis Kit (Thermo Fisher Scientific) according to the manufacturer’s protocol. qPCR was then performed on a CFX96^TM^ Real-Time System instrument (Biorad) by using the pre-amplified cDNA for the target genes. qPCR analysis was performed using Power-Up SYBR green mastermix (Thermo Fisher Scientific). Relative gene expression was calculated as previously described (Allen et al., 2017). In brief, *Rpl19* were used as housekeeping genes to generate ΔCt. ΔCt values from untreated mice were used as reference for the treatment groups to generate ΔΔCt, and relative gene expression was calculated as 2(-ΔΔCt). All reactions were run in duplicate.

#### Immunostaining

For the preparation of frozen sections, excised tumors and lymph nodes were fixed in 2% paraformaldehyde (PFA) at 4°C overnight followed by 30% sucrose at 4°C overnight before being embedded in OCT (Leica). For the preparation of agarose sections, tissues were fixed in 2% PFA at 4°C for 24h, then embedded in 4% low melting agarose (in PBS).

8μm thick or 100μm thick tumor tissue sections were stained with anti-MECA79 (clone MECA-79), anti-TCF1 (clone C63D), anti-CD8a (53-6.7), anti-PD-1 (polyclonal), anti-CD3(17A2), anti-B220 (RA3-6B2), anti-CD4 (RM4-5), anti-FOXP3 (FJK-16s), anti-CD31 (polyclonal; 2H8), anti-TIM3 (E9K5D). When primary antibodies were unlabeled, fluorophore-conjugated secondary antibodies were used. Images were taken with an Observer Z1 microscope (Zeiss) linked to an AxioCam MRM camera (Zeiss) with objectives of 20X magnification, or Leica Confocal SP8 with objectives of 20X or 63X magnification.

HEV quantification on tissue sections was assessed by quantifying the number of CD31^+^ MECA79^+^ tumor vessels. When specified, the HEV number was normalized to the non-necrotic tumor area. For evaluation of necrotic areas, frozen tumor sections were stained with hematoxylin-eosin (H&E). H&E sections were subsequently imaged under fluorescence conditions (577nm excitation wavelength) using an Observer Z1 microscope (Zeiss) (10x magnification) or Zeiss Axio Scan.Z1 (20X magnification), allowing for identification of necrotic regions through autofluorescent properties of necrotic cells (Allen et al., 2017). on-necrotic area was measured by subtracting the necrotic area to the total tumor area. Image analysis and quantitation were performed using ImageJ software version 2.0.0.

#### MILAN multiplex immunohistochemistry

##### Tissue staining

Multiple Iteractive Labeling by Antibody Neodeposition (MILAN) immunohistochemistry was performed according to a previously published method (Bolognesi et al., 2017; Bosisio et al., 2020; Giorgio et al., 2021). Tissue sections (5 μm) were prepared from FFPE murine MC38 samples. Following dewaxing, antigen retrieval was performed using PT link (Agilent) using 10 mM EDTA in Tris-buffer pH 8. Immunofluorescence staining was performed using Bond RX Fully Automated Research Stainer (Leica Biosystems) with the following primary antibodies: anti-CD31 (polyclonal); TIM3 (clone EPR22241); CD8 (clone 4SM15); PD1 (D7D5W), GrzB (polyclonal); Ki67 (clone D3B5); MECA79 (clone MECA79); TCF1 (clone C63D9). The sections were incubated for 4 hours with the primary antibodies, washed and then incubated for 30 minutes with secondary antibodies. To amplify the signal, negative-isotype controls were added for 30 min followed by a second incubation with the same secondary antibody for 30 min after three washing steps. A coverslip was placed into the slides with medium containing 4,6-diamidino-2-phenylindole (DAPI) and were scanned using a Zeiss Axio Scan Z.1 (Zeiss) at 10X magnification. The coverslips were removed after 30 min soaking in washing buffer. Stripping of the antibodies was performed in a buffer containing 1% SDS and β-mercaptoethanol for 30 min at 56 °C. After. The staining procedure was repeated for several rounds until all markers were stained and scanned.

##### Image processing

Sequential rounds were aligned using an FFT-based registration method (Matungka et al., 2009) in the DAPI channel of every round. Cell objects were defined using Stardist (Schmidt et al., 2018) in the DAPI channel for the first round. Signal autofluorescence was removed by applying a weighted subtraction of the reference round. A mask for every marker (CD8, CD31, MECA79, PD1, TIM3, TCF1, Ki67, GrzB) was obtained by applying a high-pass filter with an adaptive threshold. CD8^+^ objects were identified as the segmented cells with 50% or more overlap with the CD8 mask. The following structures of interest were defined using these masks: TCF1^+^ PD1^+^ TIM3^-^= CD8^+^ pT_EX_; TCF1^-^ PD1^+^ TIM3^+^= CD8^+^ tT_EX_; TCF1^+^ PD1^-^ TIM3^-^= CD8^+^ TCF1 cells; CD31^+^ MECA79^-^= blood vessels; CD31^+^ MECA79^+^=HEVs. The rest of CD8 T cells were identified as “not otherwise specified” or NOS for short.

##### Data analysis

By segmenting the tissue in areas of 100 sq micrometers, we then classified the tissue into different vascular areas dependent on the percentage of MECA79^+^CD31^+^ HEVs and MECA79^neg^ CD31^+^ BVs. If more than 25% of the vessels inside each area were HEVs the area was considered as HEV-high, if less as HEV-low. If CD31^+^ BVs were sparsely apparent or non-apparent, the area was considered non-vascular. Expert pathologist (Dr. Francesca Maria Bosisio) annotated the tissue sections in three different areas: tumor-bulk, tumor-edge, and non-tumor. The percentage of the defined CD8 T cell subsets (pT_EX_ vs pT_EX_ vs TCF1^+^PD1^-^ vs nos) as well as their proliferation status (Ki67^+^ vs Ki67^-^) and cytotoxicity (average levels of GrzB) was measured in the different tumoral and vascular areas. The statistical test used was the Wilcoxon signed-rank test. Adjustment for multiple tests was performed using the false-discovery-rate (FDR) method.

#### Single-cell RNA Sequencing

##### 10X Genomics

The single cell suspensions were converted to barcoded scRNA-seq libraries using the Chromium Single Cell 3’ Library, Gel Bead & Multiplex Kit and Chip Kit (10x Genomics), aiming for 6,000 cells per library. Samples were processed using kits pertaining to V2 barcoding chemistry of 10x Genomics. Single samples were always processed in a single well of a PCR plate, allowing all cells from a sample to be treated with the same master mix and in the same reaction vessel. For each experiment, all samples were processed in parallel in the same thermal cycler. Libraries were sequenced on an Illumina HiSeq4000, and mapped to the human genome (buildGRCh38) or to the mouse genome (build mm10) using CellRanger software (10x Genomics, version 3.0.2).

##### SmartSeq2

CD45^-^CD31^+^MECA79^+^ cells (TU_HEV and LN_HEV), or CD45^-^CD31^+^MECA79^-^ (TU_EC) cells were sorted (BD FACSAria III) in 96 well plates (VWR, DNase, RNase free) containing 2 μL of lysis buffer (0.2% Triton X-100, 4U of RNase inhibitor, Takara) per well. Plates were properly sealed and spun down at 2000 g for 1 min before storing at −80°C. Whole transcriptome amplification was performed with a modified SMART-seq2 protocol as described previously (Picelli et al., 2014), using 23 instead of 18 cycles of cDNA amplification. PCR purification was realized with a 0.8:1 ratio (ampureXP beads:DNA). Amplified cDNA quality was monitored with a high sensitivity DNA chip (Agilent) using the Bioanalyzer (Agilent). Sequencing libraries were performed using the Nextera XT kit (Illumina) as described previously (Picelli et al., 2014), using 1/4th of the recommended reagent volumes and 1/5th of input DNA with a tagmentation time of 9 min. Library quality was monitored with a high sensitivity DNA chip (Agilent) using the Bioanalyzer (Agilent). Indexing was performed with the Nextera XT index Kit V2 (A-D). Up to 4×96 single cells were pooled per sequencing lane. Samples were sequenced on the Illumina NextSeq 500 platform using 75bp single-end reads.

#### Single-Cell Transcriptomics Analysis

##### Quality control, data cleaning and normalization

Raw gene expression matrices generated per sample were analyzed with the Seurat3 package in R (Stuart et al., 2019). For Smarq-Seq2 datasets (Figure 1), 3 samples (TU ECs, TU HEVs and LN HEVs) were merged together and cells were filtered by nFeature_RNA (genes detected) > 2000 and percent.mt (percentage of mitochondria genes) < 20. For mouse 10X datasets (Figure 2), 5 samples (E0771 UT, E0771 DPAg, PyMT UT, PyMT DPAg, PyMT DPAg+anti-IFNγ) were merged together and cells were filtered by nCount_RNA (unique molecular identifiers, UMIs) > 5000 and percent.mt < 10. For human 10X datasets (Figure 2), 55 samples (from 28 patients) of VIB Grand Challenges Program (GCP) were merged and cells were filtered by nFeature_RNA > 1000 and percent.mt < 15. After filtering cells, log-normalization was performed using default NormalizeData function in Seurat. For 10X datasets (mouse and human), in silico EC selection was done by using basic Seurat pipeline of cell clustering with default parameters, followed by EC-related cluster annotation based on canonical markers, including Pecam1/PECAM1 and Cdh5/CDH5 (ECs), Prox1/PROX1 (lymphatic ECs) and Pdgfrb/PDGFRB (pericytes) to discriminate ECs from contaminating cells. The following analyses were done on ECs only.

##### Dimension Reduction, Sample Integration, Clustering and Visualization

High variable genes were selected by FindVariableFeatures and auto-scaled by ScaleData function, and a principal component analysis (PCA) was performed for all datasets using default RunPCA function in Seurat package. For the Smarq-Seq2 datasets (Figure 1), t-distributed stochastic neighbor embedding (Muntner et al., 2016) was performed using RunTSNE function in Seurat, using PCA dimensions 1 to 5, and clustering based on the sample origin (TU EC, TU HEV and LN HEV). For mouse 10X datasets (Figure 2), batch effect correction of each sample was done using Harmony algorithm (Korsunsky et al., 2019) based on PCA space, followed by uniform manifold approximation and projection (UMAP) using RunUMAP step (dims = 10) for data visualization, and FindNeighbors followed by FindClusters function (dims = 10, resolution = 0.55) in Seurat package for unsupervised clustering. For the human breast cancer datasets (Bassez et al., 2021) (Figure 3), batch effect correction of each sample was done using Harmony algorithm, followed by UMAP (dims = 5) for data visualization, and unsupervised clustering (dims = 6, resolution = 0.55) in Seurat. EC subtypes were identified mainly based on marker genes reported in the literature (Goveia et al., 2020).

##### SCENIC and Gene Set Enrichment Analysis

To carry out transcription factor network inference, SCENIC workflow was performed using Nextflow pipeline (Aibar et al., 2017) and regulon activity of each cell was evaluated using AUCell score with Bioconductor package AUCell. For functional/pathway analysis, gene set lists were collected from databases including Gene Ontology (GO), Reactome and Hallmark geneset of MSigDB database (http://www.gsea-msigdb.org/). For Gene Set Enrichment Analysis (GSEA), the enrichment of given gene sets of each cell was evaluated using AUCell package as well.

##### Trajectory Analysis

For mouse 10X datasets, we first integrated the datasets by tumor model and anti-IFNγ treatment using Harmony algorithm, and used Python package Palantir (Setty et al., 2019) to calculate diffusion map based on Harmony space, then diffusion components and visualized the data by tSNE. To predict the differentiation direction, we conducted Velocyto pipeline (La Manno et al., 2018) using the *.bam file and barcode information generated by CellRanger, and used ScVelo in Python (Bergen et al., 2020) for better visualization. The differential potential of each cell was predicted using either CytoTRACE (Gulati et al., 2020) online tool or Palantir algorithm.

##### Differential Expression Analysis and Data Visualization

Differentially expressed genes (DEGs) were identified by Wilcoxon Rank Sum test using FindMarkers function in Seurat. Gene expression levels or gene sets enrichment (AUCell score) were shown in t-score or z-score for heatmaps or waterfall plots. UMAP or tSNE plots were done using DimPlot or FeaturePlot functions in Seurat. Heatmaps, modified stacked violin plots, waterfall plots, bar plots of cluster proportion, and GSEA plots were generated using customized codes in R, and these functions were integrated into R package “SeuratExtend” which is available on Github (https://github.com/huayc09/SeuratExtend).

#### RNAscope *In Situ* Hybridization and Quantification

For the preparation of Formalin-Fixed Paraffin Embedded (FFPE) sections, tumors were excised and fixed in 10% neutral buffered formalin (Sigma-Merck) for 20h at +4℃, dehydrated and embedded in paraffin. 5mm thick FFPE sections were subjected to RNAscope in situ hybridization using the RNAscope Multiplex Fluorescent v2 assay (ACDBio) according to the manufacturer’s instructions (USM-323100 Multiplex Fluorescent v2 User Manual, MK- 5150_TN Multiplex Fluorescent V2 with ICW and 4-Plex Ancillary Kit for Manual multiplex fluorescent kit). Briefly, after deparaffinization, the slides were incubated with hydrogen peroxide for 10 min at RT. After washing, manual target retrieval was performed followed by incubation with primary antibody at 4℃ ON. Protease Plus was applied followed by hybridization with the RNAscope probes and the RNAscope 3-plex Positive (low expression Polr2a, medium expression PPIB, and high expression UBC) and Negative Control Probes. A negative control probe targeting a bacterial gene was used to assess background. Slides were then processed according to the RNAscope Multiplex Fluorescent v2 protocol (Hybridization, Amplification, and Signal Development), prior to secondary antibody incubation. Images were acquired using Vectra® Polaris™ Automated Quantitative Pathology Imaging System. For quantification, QuPath software was used to auto-detect cells and subcellular particles following the tutorial on ACDBio website (2021 Mar 30 - ACD Support Webinar: Visualization and Analysis of RNAscope™ Results using QuPath).

### QUANTIFICATION AND STATISTICAL ANALYSIS

Data entry and all analyses were performed in a blinded fashion. In fig5B, the tumor size measurement has been taken at the indicated timepoints +/- one day. In fig4B and figS4C and I, the label 8D off is referred at samples collected 8-9 days after treatment cessation; the label 18D off is referred at samples collected 14-20 days after treatment cessation.

Bar graphs show mean values ± SEM. Unpaired Student’s t-test (two-tailed) (Mann-Withney U test) was used for the comparison of two groups. The Wilcoxon test was used to compare two paired groups. Kruskal-Wallis test or 2way ANOVA was used for comparing > 2 groups as indicated in the corresponding legends. Statistics were indicated only when significant. p values < 0.05 were considered significant (*: p < 0.05; **: p < 0.01; ***: p < 0.001; ****: p < 0.0001). All statistical analyses were performed using GraphPad Prism software (Version 9.2) or ggpubr package in R.

## Supplemental Information

See Pdf file

## Acknowledgements

The authors thank J. Browning for his kind gift of mouse LT-b receptor-Ig Fusion protein; K. Feyen (VIB-KU Leuven) and N. Dupont (VIB-KU Leuven) for their technical assistance, P. Zhao (VIB-KU Leuven) for assistance with scRNA-seq analysis, the KU Leuven FACS core and O. Burton (University of Cambridge) for assistance with flow cytometry, the VIB histology core for assistance with histology, and S. Vlayen (VIB-KU Leuven) for the administrative and logistical support. This work was supported by grants from the Flemish government FWO (G0A0818N to GB), the National Institute of Health NIH/NCI (R01CA201537 to GB) and the Kom op tegen Kanker, Emmanuel van der Schueren starter fellowship (2018/11321/2822 to YH).

## Author Contributions

Y.H., G.V., S.S, G.B., designed experiments ; Y.H., G.V., E.A., D.N., S.J., M.D., performed all experiments unless specified; A.S., S.D., T.H., G.F.,P.M., D.L., A.L., provided and generated critical reagents, software and input; A.A, F.M.B., analyzed MILAN multiplex immunohistochemistry; G.B., S.S., F.R., conceptually planned and supervised the study. G.B wrote, Y.H., G.V., S.S., F.R. edited the manuscript.

## References

Ager, A. (2017). High Endothelial Venules and Other Blood Vessels: Critical Regulators of Lymphoid Organ Development and Function. Front Immunol 8, 45.

Aibar, S., Gonzalez-Blas, C.B., Moerman, T., Huynh-Thu, V.A., Imrichova, H., Hulselmans, G., Rambow, F., Marine, J.C., Geurts, P., Aerts, J., et al. (2017). SCENIC: single-cell regulatory network inference and clustering. Nat Methods 14, 1083–1086.

Allen, E., Jabouille, A., Rivera, L.B., Lodewijckx, I., Missiaen, R., Steri, V., Feyen, K., Tawney, J., Hanahan, D., Michael, I.P., et al. (2017). Combined antiangiogenic and anti-PD-L1 therapy stimulates tumor immunity through HEV formation. Science Translational Medicine 9.

Anandappa, A.J., Wu, C.J., and Ott, P.A. (2020). Directing Traffic: How to Effectively Drive T Cells into Tumors. Cancer Discov 10, 185–197.

Ayers, M., Lunceford, J., Nebozhyn, M., Murphy, E., Loboda, A., Kaufman, D.R., Albright, A., Cheng, J.D., Kang, S.P., Shankaran, V., et al. (2017). IFN-gamma-related mRNA profile predicts clinical response to PD-1 blockade. J Clin Invest 127, 2930–2940.

Bassez, A., Vos, H., Van Dyck, L., Floris, G., Arijs, I., Desmedt, C., Boeckx, B., Vanden Bempt, M., Nevelsteen, I., Lambein, K., et al. (2021). A single-cell map of intratumoral changes during anti-PD1 treatment of patients with breast cancer. Nat Med 27, 820–832.

Bergen, V., Lange, M., Peidli, S., Wolf, F.A., and Theis, F.J. (2020). Generalizing RNA velocity to transient cell states through dynamical modeling. Nat Biotechnol 38, 1408–1414.

Bolognesi, M.M., Manzoni, M., Scalia, C.R., Zannella, S., Bosisio, F.M., Faretta, M., and Cattoretti, G. (2017). Multiplex Staining by Sequential Immunostaining and Antibody Removal on Routine Tissue Sections. J Histochem Cytochem 65, 431–444.

Bosisio, F.M., Antoranz, A., van Herck, Y., Bolognesi, M.M., Marcelis, L., Chinello, C., Wouters, J., Magni, F., Alexopoulos, L., Stas, M., et al. (2020). Functional heterogeneity of lymphocytic patterns in primary melanoma dissected through single-cell multiplexing. Elife 9.

Browaeys, R., Saelens, W., and Saeys, Y. (2020). NicheNet: modeling intercellular communication by linking ligands to target genes. Nat Methods 17, 159–162.

Browning, J.L., Allaire, N., Ngam-Ek, A., Notidis, E., Hunt, J., Perrin, S., and Fava, R.A. (2005). Lymphotoxin-beta receptor signaling is required for the homeostatic control of HEV differentiation and function. Immunity 23, 539–550.

Browning, J.L., Sizing, I.D., Lawton, P., Bourdon, P.R., Rennert, P.D., Majeau, G.R., Ambrose, C.M., Hession, C., Miatkowski, K., Griffiths, D.A., et al. (1997). Characterization of lymphotoxin-alpha beta complexes on the surface of mouse lymphocytes. J Immunol 159, 3288–3298.

Brulois, K., Rajaraman, A., Szade, A., Nordling, S., Bogoslowski, A., Dermadi, D., Rahman, M., Kiefel, H., O’Hara, E., Koning, J.J., et al. (2020). A molecular map of murine lymph node blood vascular endothelium at single cell resolution. Nat Commun 11, 3798.

Butcher, L.M., Ito, M., Brimpari, M., Morris, T.J., Soares, F.A.C., Ahrlund-Richter, L., Carey, N., Vallier, L., Ferguson-Smith, A.C., and Beck, S. (2016). Non-CG DNA methylation is a biomarker for assessing endodermal differentiation capacity in pluripotent stem cells. Nat Commun 7, 10458.

Colbeck, E.J., Ager, A., Gallimore, A., and Jones, G.W. (2017). Tertiary Lymphoid Structures in Cancer: Drivers of Antitumor Immunity, Immunosuppression, or Bystander Sentinels in Disease? Front Immunol 8, 1830.

Gago da Graca, C., van Baarsen, L.G.M., and Mebius, R.E. (2021). Tertiary Lymphoid Structures: Diversity in Their Development, Composition, and Role. J Immunol 206, 273–281.

Giorgio, C., Francesca Maria, B., Lukas, M., and Maddalena Maria, B. (2021). Protocol Exchange.

Girard, J.P., Moussion, C., and Forster, R. (2012). HEVs, lymphatics and homeostatic immune cell trafficking in lymph nodes. Nat Rev Immunol 12, 762–773.

Goveia, J., Rohlenova, K., Taverna, F., Treps, L., Conradi, L.C., Pircher, A., Geldhof, V., de Rooij, L., Kalucka, J., Sokol, L., et al. (2020). An Integrated Gene Expression Landscape Profiling Approach to Identify Lung Tumor Endothelial Cell Heterogeneity and Angiogenic Candidates. Cancer Cell 37, 421.

Gulati, G.S., Sikandar, S.S., Wesche, D.J., Manjunath, A., Bharadwaj, A., Berger, M.J., Ilagan, F., Kuo, A.H., Hsieh, R.W., Cai, S., et al. (2020). Single-cell transcriptional diversity is a hallmark of developmental potential. Science 367, 405–411.

He, B., Jabouille, A., Steri, V., Johansson-Percival, A., Michael, I.P., Kotamraju, V.R., Junckerstorff, R., Nowak, A.K., Hamzah, J., Lee, G., et al. (2018). Vascular targeting of LIGHT normalizes blood vessels in primary brain cancer and induces intratumoural high endothelial venules. J Pathol 245, 209–221.

Hindley, J.P., Jones, E., Smart, K., Bridgeman, H., Lauder, S.N., Ondondo, B., Cutting, S., Ladell, K., Wynn, K.K., Withers, D., et al. (2012). T-cell trafficking facilitated by high endothelial venules is required for tumor control after regulatory T-cell depletion. Cancer Res 72, 5473–5482.

Hodi, F.S., O’Day, S.J., McDermott, D.F., Weber, R.W., Sosman, J.A., Haanen, J.B., Gonzalez, R., Robert, C., Schadendorf, D., Hassel, J.C., et al. (2010). Improved survival with ipilimumab in patients with metastatic melanoma. N Engl J Med 363, 711–723.

Homeister, J.W., Thall, A.D., Petryniak, B., Maly, P., Rogers, C.E., Smith, P.L., Kelly, R.J., Gersten, K.M., Askari, S.W., Cheng, G., et al. (2001). The alpha(1,3)fucosyltransferases FucT-IV and FucT-VII exert collaborative control over selectin-dependent leukocyte recruitment and lymphocyte homing. Immunity 15, 115–126.

Jain, R.K. (2001). Normalizing tumor vasculature with anti-angiogenic therapy: a new paradigm for combination therapy. Nat Med 7, 987–989.

Jansen, C.S., Prokhnevska, N., Master, V.A., Sanda, M.G., Carlisle, J.W., Bilen, M.A., Cardenas, M., Wilkinson, S., Lake, R., Sowalsky, A.G., et al. (2019). An intra-tumoral niche maintains and differentiates stem-like CD8 T cells. Nature 576, 465–470.

Jeucken, K.C.M., Koning, J.J., Mebius, R.E., and Tas, S.W. (2019). The Role of Endothelial Cells and TNF-Receptor Superfamily Members in Lymphoid Organogenesis and Function During Health and Inflammation. Front Immunol 10, 2700.

Johansson-Percival, A., and Ganss, R. (2021). Therapeutic Induction of Tertiary Lymphoid Structures in Cancer Through Stromal Remodeling. Front Immunol 12, 674375.

Johansson-Percival, A., He, B., Li, Z.J., Kjellen, A., Russell, K., Li, J., Larma, I., and Ganss, R. (2017). De novo induction of intratumoral lymphoid structures and vessel normalization enhances immunotherapy in resistant tumors. Nat Immunol 18, 1207–1217.

Johansson-Percival, A., Li, Z.J., Lakhiani, D.D., He, B., Wang, X., Hamzah, J., and Ganss, R. (2015). Intratumoral LIGHT Restores Pericyte Contractile Properties and Vessel Integrity. Cell Rep 13, 2687–2698.

Joshi, N.S., Akama-Garren, E.H., Lu, Y., Lee, D.Y., Chang, G.P., Li, A., DuPage, M., Tammela, T., Kerper, N.R., Farago, A.F., et al. (2015). Regulatory T Cells in Tumor-Associated Tertiary Lymphoid Structures Suppress Anti-tumor T Cell Responses. Immunity 43, 579–590.

Kawashima, H., Petryniak, B., Hiraoka, N., Mitoma, J., Huckaby, V., Nakayama, J., Uchimura, K., Kadomatsu, K., Muramatsu, T., Lowe, J.B., et al. (2005). N-acetylglucosamine-6-O-sulfotransferases 1 and 2 cooperatively control lymphocyte homing through L-selectin ligand biosynthesis in high endothelial venules. Nat Immunol 6, 1096–1104.

Korsunsky, I., Millard, N., Fan, J., Slowikowski, K., Zhang, F., Wei, K., Baglaenko, Y., Brenner, M., Loh, P.R., and Raychaudhuri, S. (2019). Fast, sensitive and accurate integration of single-cell data with Harmony. Nat Methods 16, 1289–1296.

Kurtulus, S., Madi, A., Escobar, G., Klapholz, M., Nyman, J., Christian, E., Pawlak, M., Dionne, D., Xia, J., Rozenblatt-Rosen, O., et al. (2019). Checkpoint Blockade Immunotherapy Induces Dynamic Changes in PD-1(-)CD8(+) Tumor-Infiltrating T Cells. Immunity 50, 181–194 e186.

La Manno, G., Soldatov, R., Zeisel, A., Braun, E., Hochgerner, H., Petukhov, V., Lidschreiber, K., Kastriti, M.E., Lonnerberg, P., Furlan, A., et al. (2018). RNA velocity of single cells. Nature 560, 494–498.

Lee, M., Kiefel, H., LaJevic, M.D., Macauley, M.S., Kawashima, H., O’Hara, E., Pan, J., Paulson, J.C., and Butcher, E.C. (2014). Transcriptional programs of lymphoid tissue capillary and high endothelium reveal control mechanisms for lymphocyte homing. Nat Immunol 15, 982–995.

Lingscheid, T., Kurth, F., Clerinx, J., Marocco, S., Trevino, B., Schunk, M., Munoz, J., Gjorup, I.E., Jelinek, T., Develoux, M., et al. (2017). Schistosomiasis in European Travelers and Migrants: Analysis of 14 Years TropNet Surveillance Data. Am J Trop Med Hyg 97, 567–574.

Maly, P., Thall, A., Petryniak, B., Rogers, C.E., Smith, P.L., Marks, R.M., Kelly, R.J., Gersten, K.M., Cheng, G., Saunders, T.L., et al. (1996). The alpha(1,3)fucosyltransferase Fuc-TVII controls leukocyte trafficking through an essential role in L-, E-, and P-selectin ligand biosynthesis. Cell 86, 643–653.

Manavski, Y., Lucas, T., Glaser, S.F., Dorsheimer, L., Gunther, S., Braun, T., Rieger, M.A., Zeiher, A.M., Boon, R.A., and Dimmeler, S. (2018). Clonal Expansion of Endothelial Cells Contributes to Ischemia-Induced Neovascularization. Circ Res 122, 670–677.

Martin, J.D., Seano, G., and Jain, R.K. (2019). Normalizing Function of Tumor Vessels: Progress, Opportunities, and Challenges. Annu Rev Physiol 81, 505–534.

Martinet, L., Filleron, T., Le Guellec, S., Rochaix, P., Garrido, I., and Girard, J.P. (2013). High endothelial venule blood vessels for tumor-infiltrating lymphocytes are associated with lymphotoxin beta-producing dendritic cells in human breast cancer. J Immunol 191, 2001–2008.

Martinet, L., Garrido, I., Filleron, T., Le Guellec, S., Bellard, E., Fournie, J.J., Rochaix, P., and Girard, J.P. (2011). Human solid tumors contain high endothelial venules: association with T- and B-lymphocyte infiltration and favorable prognosis in breast cancer. Cancer Res 71, 5678–5687.

Matungka, R., Zheng, Y.F., and Ewing, R.L. (2009). Image registration using adaptive polar transform. IEEE Trans Image Process 18, 2340–2354.

Mazzone, M., and Bergers, G. (2019). Regulation of Blood and Lymphatic Vessels by Immune Cells in Tumors and Metastasis. Annu Rev Physiol 81, 535–560.

Melrose, J., Tsurushita, N., Liu, G., and Berg, E.L. (1998). IFN-gamma inhibits activation-induced expression of E- and P-selectin on endothelial cells. J Immunol 161, 2457–2464.

Miller, B.C., Sen, D.R., Al Abosy, R., Bi, K., Virkud, Y.V., LaFleur, M.W., Yates, K.B., Lako, A., Felt, K., Naik, G.S., et al. (2019). Subsets of exhausted CD8(+) T cells differentially mediate tumor control and respond to checkpoint blockade. Nat Immunol 20, 326–336.

Mondor, I., Jorquera, A., Sene, C., Adriouch, S., Adams, R.H., Zhou, B., Wienert, S., Klauschen, F., and Bajenoff, M. (2016). Clonal Proliferation and Stochastic Pruning Orchestrate Lymph Node Vasculature Remodeling. Immunity 45, 877–888.

Motz, G.T., Santoro, S.P., Wang, L.P., Garrabrant, T., Lastra, R.R., Hagemann, I.S., Lal, P., Feldman, M.D., Benencia, F., and Coukos, G. (2014). Tumor endothelium FasL establishes a selective immune barrier promoting tolerance in tumors. Nat Med 20, 607–615.

Moussion, C., and Girard, J.P. (2011). Dendritic cells control lymphocyte entry to lymph nodes through high endothelial venules. Nature 479, 542–546.

Muntner, P., Becker, R.C., Calhoun, D., Chen, D., Cowley, A.W., Jr., Flynn, J.T., Grobe, J.L., Kidambi, S., Kotchen, T.A., Lackland, D.T., et al. (2016). Introduction to the American Heart Association’s Hypertension Strategically Focused Research Network. Hypertension 67, 674–680.

Onder, L., Danuser, R., Scandella, E., Firner, S., Chai, Q., Hehlgans, T., Stein, J.V., and Ludewig, B. (2013). Endothelial cell-specific lymphotoxin-beta receptor signaling is critical for lymph node and high endothelial venule formation. J Exp Med 210, 465–473.

Peske, J.D., Thompson, E.D., Gemta, L., Baylis, R.A., Fu, Y.X., and Engelhard, V.H. (2015). Effector lymphocyte-induced lymph node-like vasculature enables naive T-cell entry into tumours and enhanced anti-tumour immunity. Nat Commun 6, 7114.

Picelli, S., Faridani, O.R., Bjorklund, A.K., Winberg, G., Sagasser, S., and Sandberg, R. (2014). Full-length RNA-seq from single cells using Smart-seq2. Nature Protocols 9, 171–181.

Rivera, L.B., and Bergers, G. (2015). Intertwined regulation of angiogenesis and immunity by myeloid cells. Trends in Immunology 36, 240–249.

Rodriguez, A.B., Peske, J.D., Woods, A.N., Leick, K.M., Mauldin, I.S., Meneveau, M.O., Young, S.J., Lindsay, R.S., Melssen, M.M., Cyranowski, S., et al. (2021). Immune mechanisms orchestrate tertiary lymphoid structures in tumors via cancer-associated fibroblasts. Cell Rep 36, 109422.

Rosen, S.D. (2004). Ligands for L-selectin: homing, inflammation, and beyond. Annu Rev Immunol 22, 129–156.

Sade-Feldman, M., Jiao, Y.J., Chen, J.H., Rooney, M.S., Barzily-Rokni, M., Eliane, J.P., Bjorgaard, S.L., Hammond, M.R., Vitzthum, H., Blackmon, S.M., et al. (2017). Resistance to checkpoint blockade therapy through inactivation of antigen presentation. Nat Commun 8, 1136.

Sautes-Fridman, C., Petitprez, F., Calderaro, J., and Fridman, W.H. (2019). Tertiary lymphoid structures in the era of cancer immunotherapy. Nat Rev Cancer 19, 307–325.

Schmidt, U., Weigert, M., Broaddus, C., and Myers, G. (2018). Cell Detection with Star-Convex Polygons (Cham: Springer International Publishing).

Schmittnaegel, M., Rigamonti, N., Kadioglu, E., Cassara, A., Wyser Rmili, C., Kiialainen, A., Kienast, Y., Mueller, H.J., Ooi, C.H., Laoui, D., et al. (2017). Dual angiopoietin-2 and VEGFA inhibition elicits antitumor immunity that is enhanced by PD-1 checkpoint blockade. Sci Transl Med 9.

Schumacher, T.N., and Schreiber, R.D. (2015). Neoantigens in cancer immunotherapy. Science 348, 69–74.

Setty, M., Kiseliovas, V., Levine, J., Gayoso, A., Mazutis, L., and Pe’er, D. (2019). Characterization of cell fate probabilities in single-cell data with Palantir. Nat Biotechnol 37, 451–460.

Siddiqui, I., Schaeuble, K., Chennupati, V., Fuertes Marraco, S.A., Calderon-Copete, S., Pais Ferreira, D., Carmona, S.J., Scarpellino, L., Gfeller, D., Pradervand, S., et al. (2019). Intratumoral Tcf1(+)PD-1(+)CD8(+) T Cells with Stem-like Properties Promote Tumor Control in Response to Vaccination and Checkpoint Blockade Immunotherapy. Immunity 50, 195–211 e110.

Spranger, S., and Gajewski, T.F. (2015). A new paradigm for tumor immune escape: beta-catenin-driven immune exclusion. J Immunother Cancer 3, 43.

Stuart, T., Butler, A., Hoffman, P., Hafemeister, C., Papalexi, E., Mauck, W.M., 3rd, Hao, Y., Stoeckius, M., Smibert, P., and Satija, R. (2019). Comprehensive Integration of Single-Cell Data. Cell 177, 1888–1902 e1821.

Terai, Y., Miyagi, R., Aibara, M., Mizoiri, S., Imai, H., Okitsu, T., Wada, A., Takahashi-Kariyazono, S., Sato, A., Tichy, H., et al. (2017). Visual adaptation in Lake Victoria cichlid fishes: depth-related variation of color and scotopic opsins in species from sand/mud bottoms. BMC Evol Biol 17, 200.

Tian, L., Goldstein, A., Wang, H., Ching Lo, H., Sun Kim, I., Welte, T., Sheng, K., Dobrolecki, L.E., Zhang, X., Putluri, N., et al. (2017). Mutual regulation of tumour vessel normalization and immunostimulatory reprogramming. Nature 544, 250–254.

Topalian, S.L., Hodi, F.S., Brahmer, J.R., Gettinger, S.N., Smith, D.C., McDermott, D.F., Powderly, J.D., Carvajal, R.D., Sosman, J.A., Atkins, M.B., et al. (2012). Safety, activity, and immune correlates of anti-PD-1 antibody in cancer. N Engl J Med 366, 2443–2454.

Tumeh, P.C., Harview, C.L., Yearley, J.H., Shintaku, I.P., Taylor, E.J., Robert, L., Chmielowski, B., Spasic, M., Henry, G., Ciobanu, V., et al. (2014). PD-1 blockade induces responses by inhibiting adaptive immune resistance. Nature 515, 568–571.

Uchimura, K., Gauguet, J.M., Singer, M.S., Tsay, D., Kannagi, R., Muramatsu, T., von Andrian, U.H., and Rosen, S.D. (2005). A major class of L-selectin ligands is eliminated in mice deficient in two sulfotransferases expressed in high endothelial venules. Nat Immunol 6, 1105–1113.

van de Pavert, S.A., and Mebius, R.E. (2010). New insights into the development of lymphoid tissues. Nat Rev Immunol 10, 664–674.

Veerman, K., Tardiveau, C., Martins, F., Coudert, J., and Girard, J.P. (2019). Single-Cell Analysis Reveals Heterogeneity of High Endothelial Venules and Different Regulation of Genes Controlling Lymphocyte Entry to Lymph Nodes. Cell Rep 26, 3116–3131 e3115.

Weibel, E.R. (2012). Fifty years of Weibel-Palade bodies: the discovery and early history of an enigmatic organelle of endothelial cells. J Thromb Haemost 10, 979–984.

Wherry, E.J., and Kurachi, M. (2015). Molecular and cellular insights into T cell exhaustion. Nat Rev Immunol 15, 486–499.

