## Supplementary Figures for "Anticancer immunotherapies transition postcapillary venules into high-endothelial venules that generate TCF1+ T lymphocyte niches through a feed-forward loop"

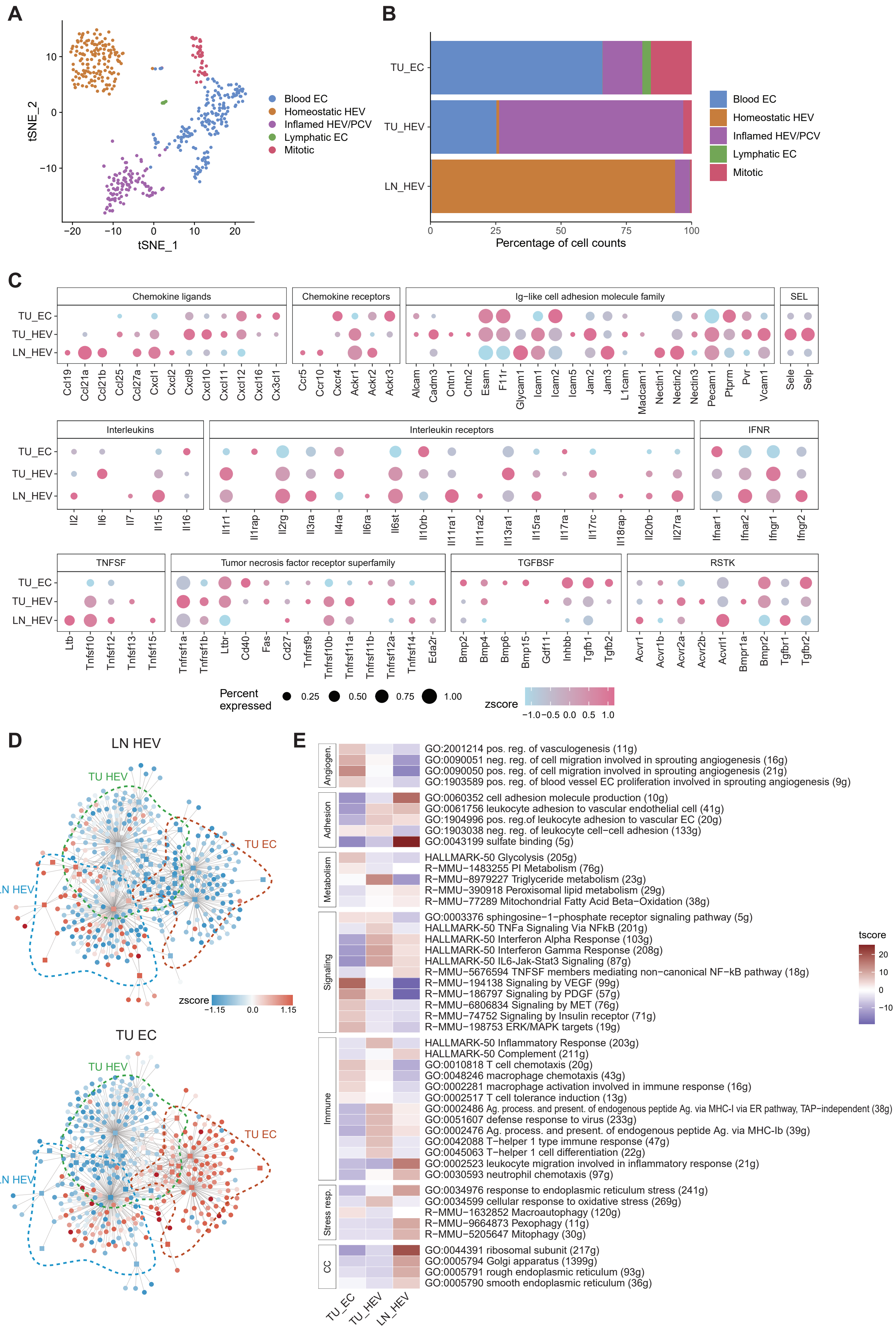

A
tSNE_2
tSNE_1
10
0
−10
−20
20
Blood EC
Homeostatic HEV
Inflamed HEV/PCV
Lymphatic EC
Mitotic
B
TU_EC
TU_HEV
LN_HEV
0
25
50
75
100
Percentage of cell counts
Blood EC
Homeostatic HEV
Inflamed HEV/PCV
Lymphatic EC
Mitotic
C
Chemokine ligands
Chemokine receptors
Ig−like cell adhesion molecule family
SEL
TU_EC
TU_HEV
LN_HEV
Ccl19 Ccl21a Ccl21b Ccl25 Ccl27a Cxcl1 Cxcl2 Cxcl9 Cxcl10 Cxcl11 Cxcl12 Cxcl16 Cx3cl1
Ccr5 Ccr10 Cxcr4 Ackr1 Ackr2 Ackr3
Alcam Cadm3 Cntn1 Cntn2 Esam F11r Glycam1 Icam1 Icam2 Icam5 Jam2 Jam3 L1cam Madcam1 Nectin1 Nectin2 Nectin3 Pecam1 Ptprm Pvr Vcam1
Sele Selp
Interleukins
Interleukin receptors
IFNR
Il2 Il6 Il7 Il15 Il16
Il1r1 Il1rap Il2rg Il3ra Il4ra Il6ra Il6st Il10rb Il11ra1 Il11ra2 Il13ra1 Il15ra Il17ra Il17rc Il18rap Il20rb Il27ra
Ifnar1 Ifnar2 Ifngr1 Ifngr2
TNFSF
Tumor necrosis factor receptor superfamily
TGFBSF
RSTK
Ltb Tnfsf10 Tnfsf12 Tnfsf13 Tnfsf15
Tnfrsf1a Tnfrsf1b Ltbr Cd40 Fas Cd27 Tnfrsf9 Tnfrsf10b Tnfrsf11a Tnfrsf11b Tnfrsf12a Tnfrsf14 Eda2r
Bmp2 Bmp4 Bmp6 Bmp15 Gdf11 Inhbb Tgfb1 Tgfb2
Acvr1 Acvr1b Acvr2a Acvr2b Acvrl1 Bmpr1a Bmpr2 Tgfbr1 Tgfbr2
Percent expressed
0.25
0.50
0.75
1.00
zscore
−1.0 −0.5 0.0 0.5 1.0
D
LN HEV
TU HEV
TU EC
LN HEV
zscore
−1.15
0.0
1.15
TU EC
TU HEV
TU EC
LN HEV
E
Angiogen.
GO:2001214 pos. reg. of vasculogenesis (11g)
GO:0090051 neg. reg. of cell migration involved in sprouting angiogenesis (16g)
GO:0090050 pos. reg. of cell migration involved in sprouting angiogenesis (21g)
GO:1903589 pos. reg. of blood vessel EC proliferation involved in sprouting angiogenesis (9g)
Adhesion
GO:0060352 cell adhesion molecule production (10g)
GO:0061756 leukocyte adhesion to vascular endothelial cell (41g)
GO:1904996 pos. reg.of leukocyte adhesion to vascular EC (20g)
GO:1903038 neg. reg. of leukocyte cell−cell adhesion (133g)
GO:0043199 sulfate binding (5g)
Metabolism
HALLMARK-50 Glycolysis (205g)
R−MMU−1483255 PI Metabolism (76g)
R−MMU−8979227 Triglyceride metabolism (23g)
R−MMU−390918 Peroxisomal lipid metabolism (29g)
R−MMU−77289 Mitochondrial Fatty Acid Beta−Oxidation (38g)
Signaling
GO:0003376 sphingosine−1−phosphate receptor signaling pathway (5g)
HALLMARK-50 TNFa Signaling Via NFkB (201g)
HALLMARK-50 Interferon Alpha Response (103g)
HALLMARK-50 Interferon Gamma Response (208g)
HALLMARK-50 IL6-Jak-Stat3 Signaling (87g)
R−MMU−5676594 TNFSF members mediating non−canonical NF−kB pathway (18g)
R−MMU−194138 Signaling by VEGF (99g)
R−MMU−186797 Signaling by PDGF (57g)
R−MMU−6806834 Signaling by MET (76g)
R−MMU−74752 Signaling by Insulin receptor (71g)
R−MMU−198753 ERK/MAPK targets (19g)
Immune
HALLMARK-50 Inflammatory Response (203g)
HALLMARK-50 Complement (211g)
GO:0010818 T cell chemotaxis (20g)
GO:0048246 macrophage chemotaxis (43g)
GO:0002281 macrophage activation involved in immune response (16g)
GO:0002517 T cell tolerance induction (13g)
GO:0002486 Ag. process. and present. of endogenous peptide Ag. via MHC-I via ER pathway, TAP−independent (38g)
GO:0051607 defense response to virus (233g)
GO:0002476 Ag. process. and present. of endogenous peptide Ag. via MHC-Ib (39g)
GO:0042088 T−helper 1 type immune response (47g)
GO:0045063 T−helper 1 cell differentiation (22g)
GO:0002523 leukocyte migration involved in inflammatory response (21g)
GO:0030593 neutrophil chemotaxis (97g)
Stress resp.
GO:0034976 response to endoplasmic reticulum stress (241g)
GO:0034599 cellular response to oxidative stress (269g)
R−MMU−1632852 Macroautophagy (120g)
R−MMU−9664873 Pexophagy (11g)
R−MMU−5205647 Mitophagy (30g)
CC
GO:0044391 ribosomal subunit (217g)
GO:0005794 Golgi apparatus (1399g)
GO:0005791 rough endoplasmic reticulum (93g)
GO:0005790 smooth endoplasmic reticulum (36g)
TU_EC
TU_HEV
LN_HEV
tscore
20
10
0
−10

**Figure S1. General characterization of TU-HEVs, LN-HEVs and TU-ECs**

(A) tSNE plot of EC transcriptomes, colored by cell type.

(B) Fraction of EC sub-groups in TU-HEV, LN-HEV and TU-EC.

(C) Expression of selected chemokines, adhesion molecules and cytokines (ligand + receptor, x axis) in three EC samples (y-axis). Dot size represents the percentage of cells in which the gene is detected. Color indicates the mean expression (in z-score).

(D) Gene regulatory network (GRN) predicted by SCENIC. node color shows the gene expression (round nodes) and regulon activity (square node) in LN-HEV and TU-EC.

(E) Heatmap showing selected pathway activities in TU-EC, TU-HEV and LN-HEV.

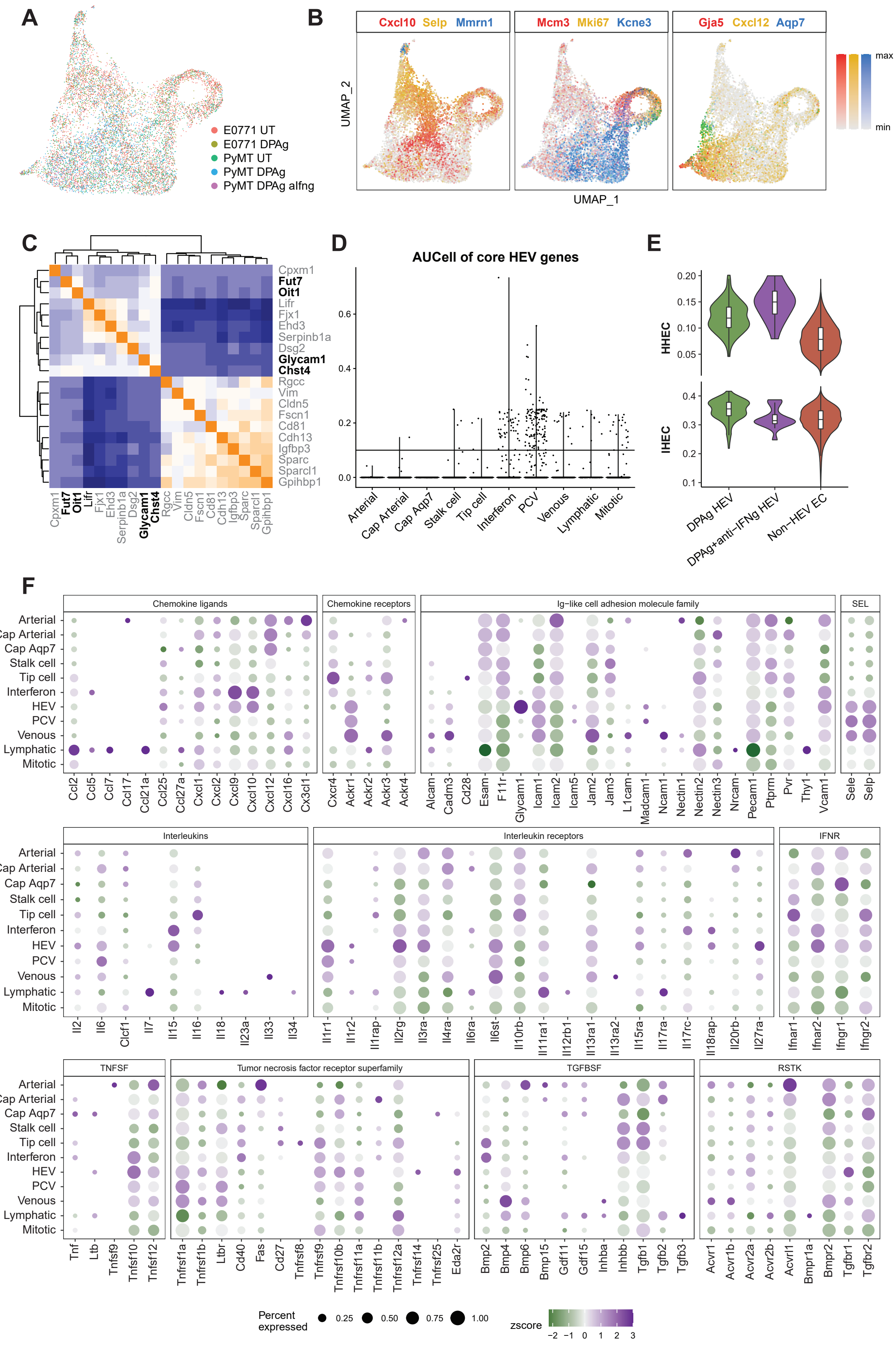
A
E0771 UT
E0771 DPAg
PyMT UT
PyMT DPAg
PyMT DPAg aIfng
B
Cxcl10 Selp Mmrn1
Mcm3 Mki67 Kcne3
Gja5 Cxcl12 Aqp7
UMAP_2
UMAP_1
max
min
C
Cpxm1
Fut7
Oit1
Lifr
Fjx1
Ehd3
Serpinb1a
Dsg2
Glycam1
Chst4
Rgcc
Vim
Cldn5
Fscn1
Cd81
Cdh13
Igfbp3
Sparc
Sparcl1
Gpihbp1
D
AUCell of core HEV genes
Arterial
Cap Arterial
Cap Aqp7
Stalk cell
Tip cell
Interferon
PCV
Venous
Lymphatic
Mitotic
E
HHEC
IHEC
DPAg HEV
DPAg+anti−IFNg HEV
Non−HEV EC
F
Chemokine ligands
Chemokine receptors
Ig−like cell adhesion molecule family
SEL
Interleukins
Interleukin receptors
IFNR
TNFSF
Tumor necrosis factor receptor superfamily
TGFBSF
RSTK
Arterial
Cap Arterial
Cap Aqp7
Stalk cell
Tip cell
Interferon
HEV
PCV
Venous
Lymphatic
Mitotic
Ccl2 Ccl5 Ccl7 Ccl17 Ccl21a Ccl25 Ccl27a Cxcl1 Cxcl2 Cxcl9 Cxcl10 Cxcl12 Cxcl16 Cx3cl1
Cxcr4 Ackr1 Ackr2 Ackr3 Ackr4
Alcam Cadm3 Cd28 Esam F11r Glycam1 Icam1 Icam2 Icam5 Jam2 Jam3 L1cam Madcam1 Ncam1 Nectin1 Nectin2 Nectin3 Nrcam Pecam1 Ptprm Pvr Thy1 Vcam1
Sele Selp
Il2 Il6 Clcf1 Il7 Il15 Il16 Il18 Il23a Il33 Il34
Il1r1 Il1r2 Il1rap Il2rg Il3ra Il4ra Il6ra Il6st Il10rb Il11ra1 Il12rb1 Il13ra1 Il13ra2 Il15ra Il17ra Il17rc Il18rap Il20rb Il27ra
Ifnar1 Ifnar2 Ifngr1 Ifngr2
Tnf Ltb Tnfsf9 Tnfsf10 Tnfsf12
Tnfrsf1a Tnfrsf1b Ltbr Cd40 Fas Cd27 Tnfrsf8 Tnfrsf9 Tnfrsf10b Tnfrsf11a Tnfrsf11b Tnfrsf12a Tnfrsf14 Tnfrsf25 Eda2r
Bmp2 Bmp4 Bmp6 Bmp15 Gdf11 Gdf15 Inhba Inhbb Tgfb1 Tgfb2 Tgfb3
Acvr1 Acvr1b Acvr2a Acvr2b Acvrl1 Bmpr1a Bmpr2 Tgfbr1 Tgfbr2
Percent expressed
0.25 0.50 0.75 1.00
zscore
−2 −1 0 1 2 3

**Figure S2. Characterization of the mouse tumor vasculature by droplet-based scRNAseq**

(A) UMAP plot, colored by samples integrated by Harmony algorithm.

(B) Expression level of representative marker genes to annotate EC subtypes.

(C) Pearson correlation matrix heatmap showing the top 10 positive and negative correlated genes with Chst4.

(D) AUCell of core HEV genes in each cell. AUCell > 0.1 are defined as HEVs.

(E) Violin plots showing the AUCell of gene signatures of HHEC (homeostatic HEC) and IHEC (inflammatory HEC) (Veerman et al., 2019) in DPAg HEVs, DPAg + anti-IFNγ HEVs and other ECs.

(F) Expression of selected chemokines, adhesion molecules and cytokines (ligand + receptor, x axis) in each EC subtype (y-axis). Dot size represents the percentage of cells in which the gene is detected. Color indicates the mean expression (in z-score).

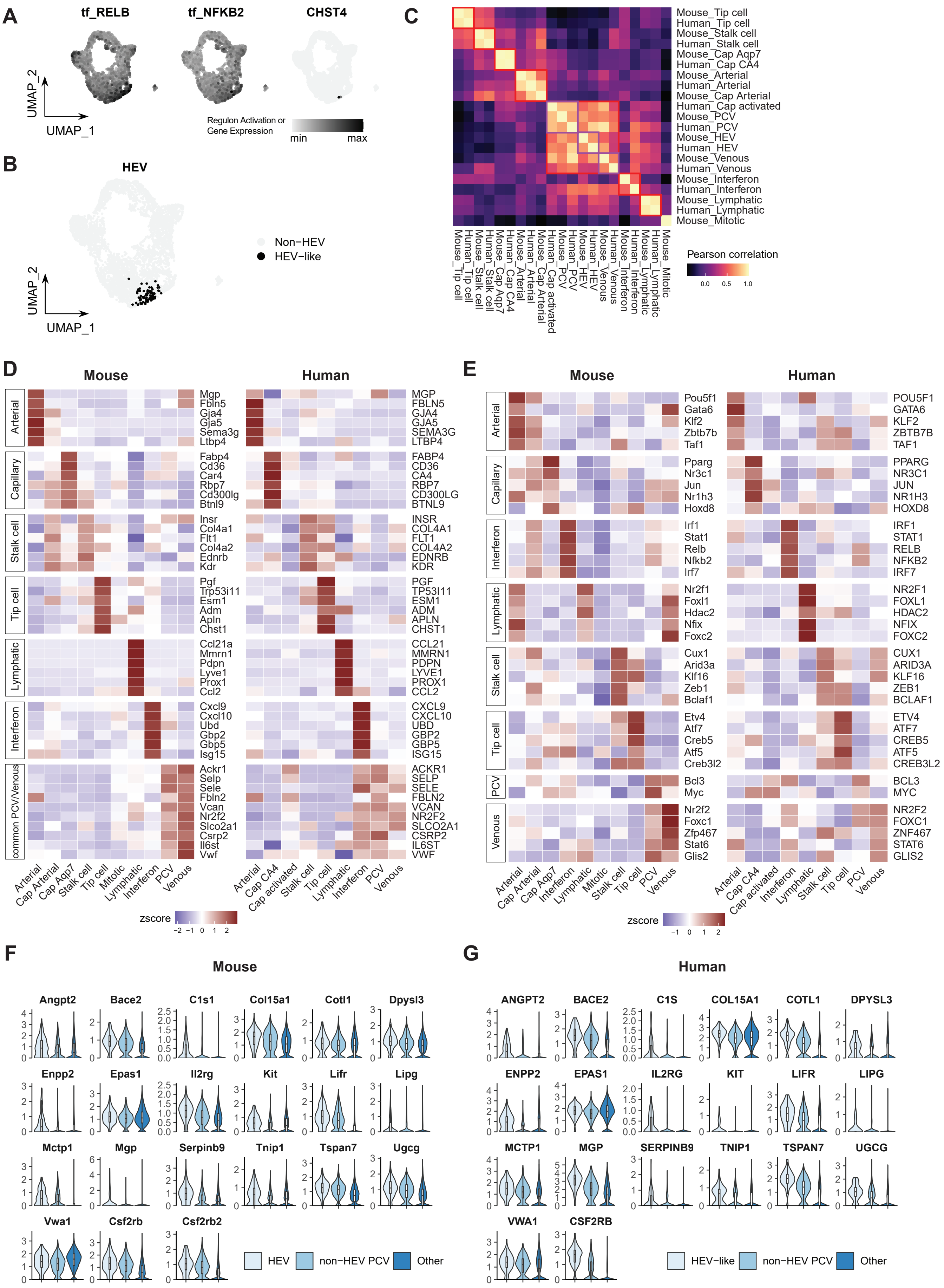
A
tf_RELB
tf_NFKB2
CHST4
UMAP_2
UMAP_1
Regulon Activation or Gene Expression
min
max
B
HEV
Non−HEV
HEV−like
C
Mouse_Tip cell
Human_Tip cell
Mouse_Stalk cell
Human_Stalk cell
Mouse_Cap Aqp7
Human_Cap CA4
Mouse_Arterial
Human_Arterial
Mouse_Cap Arterial
Human_Cap activated
Mouse_PCV
Human_PCV
Mouse_HEV
Human_HEV
Mouse_Venous
Human_Venous
Mouse_Interferon
Human_Interferon
Mouse_Lymphatic
Human_Lymphatic
Mouse_Mitotic
Pearson correlation
0.0
0.5
1.0
D
Mouse
Human
Arterial
Capillary
Stalk cell
Tip cell
Lymphatic
Interferon
common PCV/Venous
Mgp Fbln5 Gja4 Gja5 Sema3g Ltbp4
Fabp4 Cd36 Car4 Rbp7 Cd300lg Btnl9
Insr Col4a1 Flt1 Col4a2 Ednrb Kdr
Pgf Trp53i11 Esm1 Adm Apln Chst1
Ccl21a Mmrn1 Pdpn Lyve1 Prox1 Ccl2
Cxcl9 Cxcl10 Ubd Gbp2 Gbp5 Isg15
Ackr1 Selp Sele Fbln2 Vcan Nr2f2 Slco2a1 Csrp2 Il6st Vwf
MGP FBLN5 GJA4 GJA5 SEMA3G LTBP4
FABP4 CD36 CA4 RBP7 CD300LG BTNL9
INSR COL4A1 FLT1 COL4A2 EDNRB KDR
PGF TP53I11 ESM1 ADM APLN CHST1
CCL21 MMRN1 PDPN LYVE1 PROX1 CCL2
CXCL9 CXCL10 UBD GBP2 GBP5 ISG15
ACKR1 SELP SELE FBLN2 VCAN NR2F2 SLCO2A1 CSRP2 IL6ST VWF
Arterial Cap Arterial Cap Aqp7 Stalk cell Tip cell Mitotic Lymphatic Interferon PCV Venous
Arterial Cap CA4 Cap activated Stalk cell Tip cell Lymphatic Interferon PCV Venous
zscore
−2 −1 0 1 2
E
Mouse
Human
Arterial
Capillary
Interferon
Lymphatic
Stalk cell
Tip cell
PCV
Venous
Pou5f1 Gata6 Klf2 Zbtb7b Taf1
Pparg Nr3c1 Jun Nr1h3 Hoxd8
Irf1 Stat1 Relb Nfkb2 Irf7
Nr2f1 Foxl1 Hdac2 Nfix Foxc2
Cux1 Arid3a Klf16 Zeb1 Bclaf1
Etv4 Atf7 Creb5 Atf5 Creb3l2
Bcl3 Myc
Nr2f2 Foxc1 Zfp467 Stat6 Glis2
POU5F1 GATA6 KLF2 ZBTB7B TAF1
PPARG NR3C1 JUN NR1H3 HOXD8
IRF1 STAT1 RELB NFKB2 IRF7
NR2F1 FOXL1 HDAC2 NFIX FOXC2
CUX1 ARID3A KLF16 ZEB1 BCLAF1
ETV4 ATF7 CREB5 ATF5 CREB3L2
BCL3 MYC
NR2F2 FOXC1 ZNF467 STAT6 GLIS2
Arterial Cap Arterial Cap Aqp7 Interferon Lymphatic Mitotic Stalk cell Tip cell PCV Venous
Arterial Cap CA4 Cap activated Interferon Lymphatic Stalk cell Tip cell PCV Venous
zscore
−1 0 1 2
F
Mouse
Angpt2 Bace2 C1s1 Col15a1 Cotl1 Dpysl3
Enpp2 Epas1 Il2rg Kit Lifr Lipg
Mctp1 Mgp Serpinb9 Tnip1 Tspan7 Ugcg
Vwa1 Csf2rb Csf2rb2
HEV
non−HEV PCV
Other
G
Human
ANGPT2 BACE2 C1S COL15A1 COTL1 DPYSL3
ENPP2 EPAS1 IL2RG KIT LIFR LIPG
MCTP1 MGP SERPINB9 TNIP1 TSPAN7 UGCG
VWA1 CSF2RB
HEV−like
non−HEV PCV
Other

**Figure S3. Characterization of the human tumor vasculature by scRNAseq**

(A and B) UMAP plots, colored by RELB/NFKB2 regulon activities predicted by SCENIC or expression of CHST4 (A), or *in silico* selected HEV cells (B).

(C) Pearson correlation of Harmony corrected principle components (PCs) exhibiting the similarity of each EC subtype in mouse and human.

(D and E) Conserved features (D) or regulons (E) of each EC subtype in mouse and human datasets. Homologous genes are shown in the same row.

(F and G) RNA expression levels of conserved TU-HEV features in mouse (F) and human (G) breast cancer datasets, comparing TU-HEVs, non-HEV PCVs and other ECs.

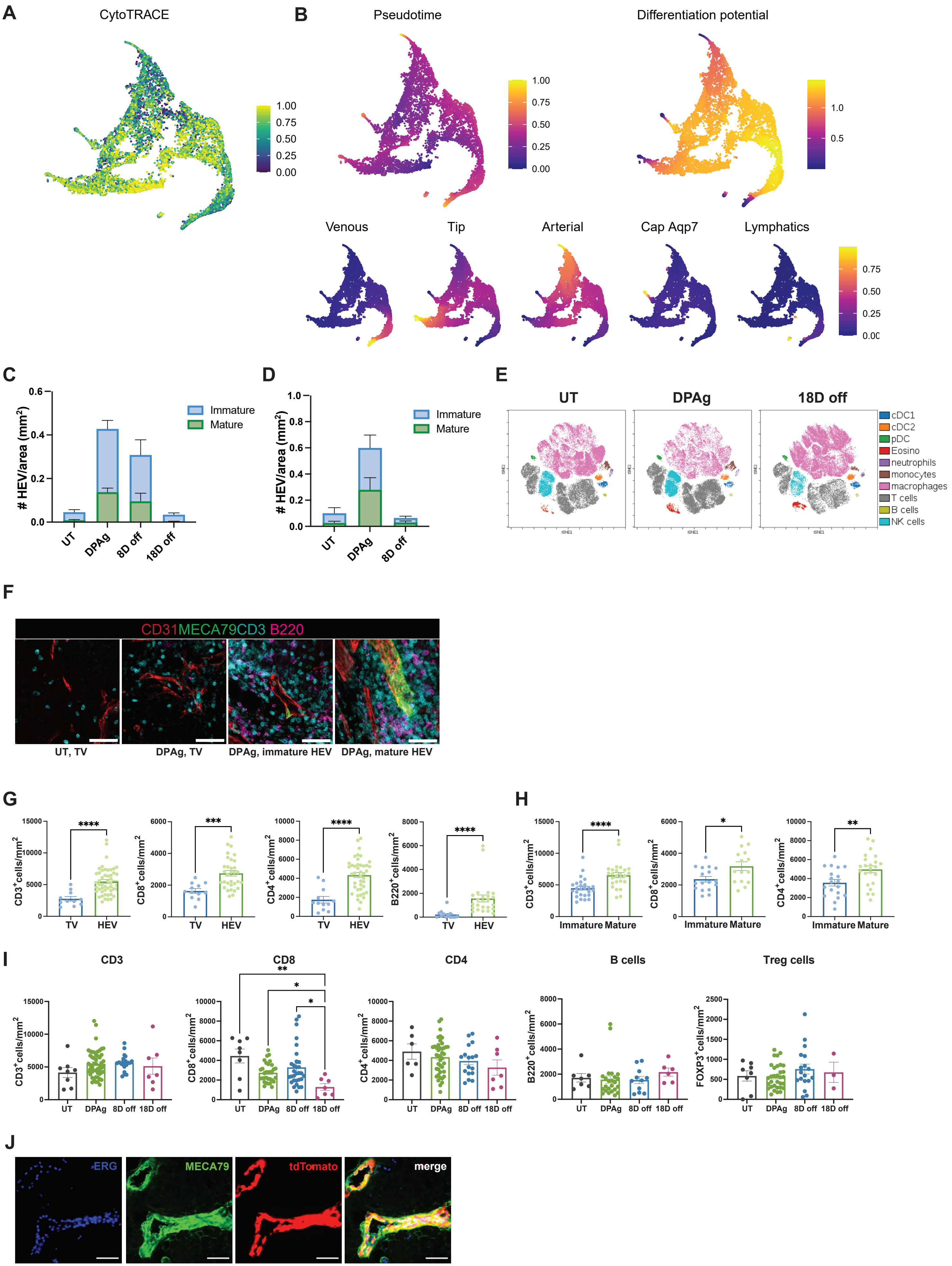
A
CytoTRACE
1.00
0.75
0.50
0.25
0.00
B
Pseudotime
Differentiation potential
1.00
0.75
0.50
0.25
0.00
1.0
0.5
Venous
Tip
Arterial
Cap Aqp7
Lymphatics
0.75
0.50
0.25
0.00
C
### HEV/area (mm²)
Immature
Mature
UT
DPAg
8D off
18D off
D
### HEV/area (mm²)
Immature
Mature
UT
DPAg
8D off
E
UT
DPAg
18D off
cDC1
cDC2
pDC
Eosino
neutrophils
monocytes
macrophages
T cells
B cells
NK cells
F
CD31MECA79CD3 B220
UT, TV
DPAg, TV
DPAg, immature HEV
DPAg, mature HEV
G
CD3⁺cells/mm²
CD8⁺cells/mm²
CD4⁺cells/mm²
B220⁺cells/mm²
TV
HEV
****
***
H
Immature
Mature
*
**
I
CD3
CD8
CD4
B cells
Treg cells
UT
DPAg
8D off
18D off
FOXP3⁺cells/mm²
J
ERG
MECA79
tdTomato
merge

**Figure S4. TU-EC metaplasia into TU-HEVs is dependent on immunotherapy-induced signals**

(A and B) Differentiation potential, predicted by CytoTRACE (A) or Palantir (B).

(C and D) Ratio of mature/immature HEV phenotypes of UT, DPAg or DPAg-stop PyMT (N tumors = 4 to 10) (C) and E0771 (N tumors = 11) (D) tumors.

(E) Representative Cytobank tSNE plot using the PyMT flow cytometry data.

(F) Representative pictures of tumor vessels (TVs) ($CD31^+MECA79^-$), immature/mature HEVs ($CD31^+MECA79^+$), B ($B220^+$) and T ($CD3^+$) lymphocytes in untreated (UT) and DPAg treated PyMT tumors. Scale bar indicates 50μm.

(G) Quantification of CD3 T cells (N = 11-51), CD8 T cells (N = 12-32), CD4 T cells (N = 12-42), and B cells (N = 13-24) $50 μm^2$ around HEVs or TVs in DPAg-treated PyMT tumors by immunofluorescent tissue staining.

(H) Quantification of CD3 T cells (N = 25-26), CD4 T cells (N = 19-23), and CD8 T cells (N = 15-17) $50 μm^2$ around mature HEVs and immature HEVs in DPAg-treated PyMT tumors by immunofluorescent tissue staining.

(I) Quantification of CD3 T cells (N = 7-51), CD8 T cells (N = 7-32), CD4 T cells (N = 6-42), B cells (N = 6-24), and Treg cells (N = 3-32) $50\ μm^2$ around HEVs from PyMT tumors by immunofluorescent tissue staining.

(J) Representative image of LN-HEVs from Chst4-tdT mice. Scale bars indicate 50 μm.

The mean and the SEM are shown. Mann-Whitney test (G and H). Krustal-Wallist test (I). Data are pooled from at least two independent experiments (C, D, G-I).

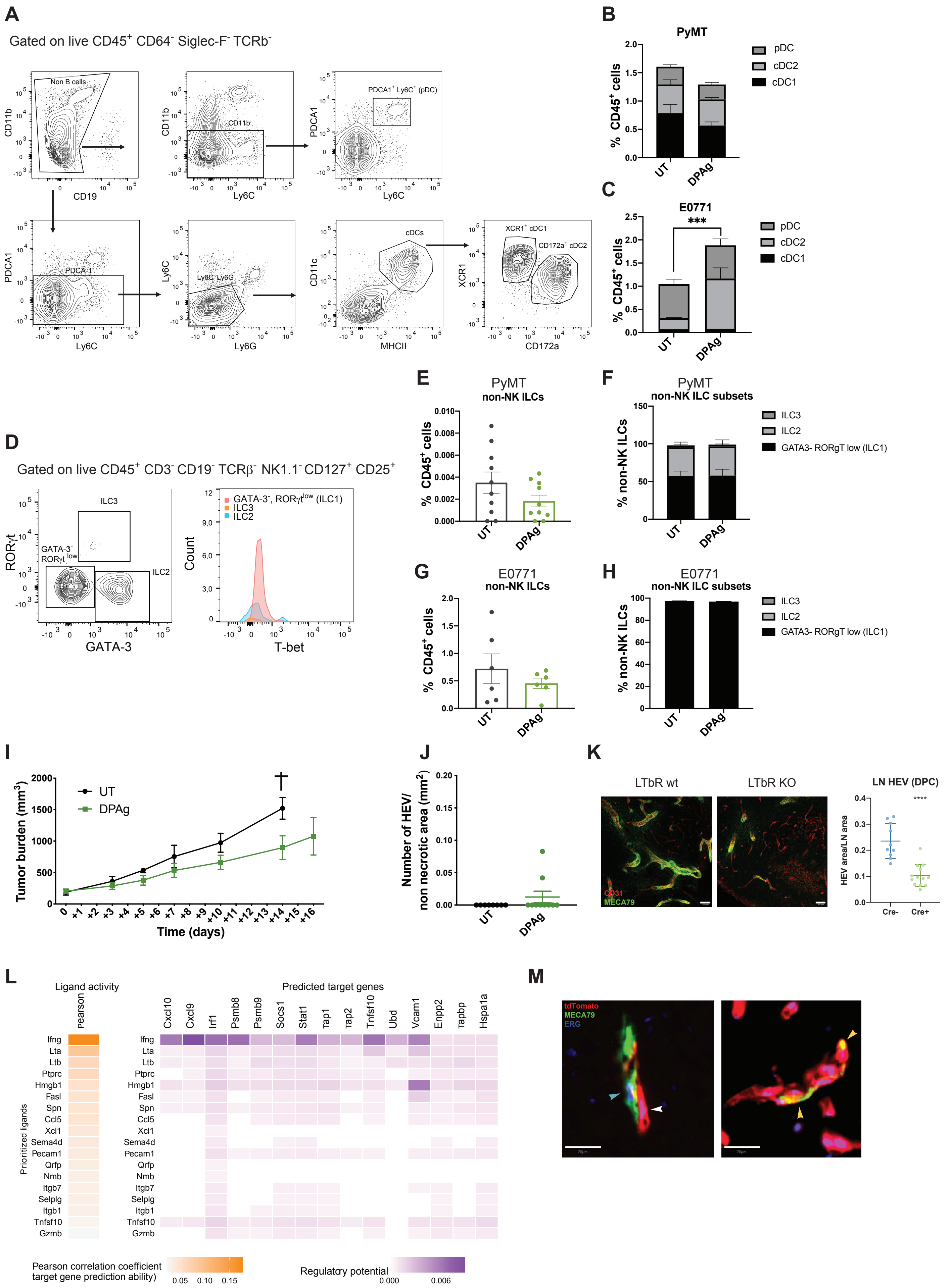

A
Gated on live CD45+ CD64- Siglec-F- TCRb-
Non B cells
CD11b
CD19
CD11b-
Ly6C
PDCA1+ Ly6C+ (pDC)
PDCA1
PDCA-1-
Ly6C-Ly6G-
Ly6G
cDCs
CD11c
MHCII
XCR1+ cDC1
CD172a+ cDC2
XCR1
CD172a
B
PyMT
% CD45+ cells
pDC
cDC2
cDC1
UT
DPAg
C
E0771
***
D
Gated on live CD45+ CD3- CD19- TCRβ- NK1.1- CD127+ CD25+
ILC3
GATA-3- RORγt low
ILC2
RORγt
GATA-3
GATA-3-, RORγtlow (ILC1)
Count
T-bet
E
PyMT
non-NK ILCs
F
PyMT
non-NK ILC subsets
% non-NK ILCs
ILC3
ILC2
GATA3- RORgT low (ILC1)
G
E0771
non-NK ILCs
H
E0771
non-NK ILC subsets
I
Tumor burden (mm3)
Time (days)
J
Number of HEV/
non necrotic area (mm2)
K
LTbR wt
LTbR KO
CD31
MECA79
LN HEV (DPC)
****
HEV area/LN area
Cre-
Cre+
L
Ligand activity
Pearson
Predicted target genes
Cxcl10
Cxcl9
Irf1
Psmb8
Psmb9
Socs1
Stat1
Tap1
Tap2
Tnfsf10
Ubd
Vcam1
Enpp2
Tapbp
Hspa1a
Prioritized ligands
Ifng
Lta
Ltb
Ptprc
Hmgb1
Fasl
Spn
Ccl5
Xcl1
Sema4d
Pecam1
Qrfp
Nmb
Itgb7
Selplg
Itgb1
Tnfsf10
Gzmb
Pearson correlation coefficient
target gene prediction ability)
0.05 0.10 0.15
Regulatory potential
0.000 0.006
M
tdTomato
MECA79
ERG

**Figure S5. CD8 T-cells and NK cells induce immunotherapy-dependent HEV formation via the LT/LTβR axis**

(A-C) Gating strategy for intratumoral XCR1$^+$ cDC1s, CD172a$^+$ cDC2s and PDCA1$^+$ pDCs by flow cytometry (A) and quantification of each DC subset in UT or DPAg treated PyMT (N = 10) (B) or E0771 (N = 6) (C) tumors.

(D) Flow cytometry gating strategy for intratumoral non-NK ILC1, ILC2 and ILC3 based on the expression of Tbet (ILC1), Gata3 (ILC2) and Rorγt (ILC3) in CD19$^-$CD3$^-$TCR$^-$NK1.1$^-$CD25$^+$ CD127$^+$ cells.

(E-H) Quantification of non-NK ILCs in PyMT (N = 10) (E) and E0771 (N = 6) (G) and of each non-NK ILC subset in PyMT (N = 10) (F) or E0771 (N = 6) (H) tumors.

(I and J) Tumor growth curve (N = 7-9) (I) and HEV density (N tumors = 8-10) (J) of UT or DPAg treated PyMT-bearing Rag1 KO mice.

(K) Representative images (left) and quantification (right) of LN-HEVs from LTβR wt (Cre$^-$) and LTβR KO (Cre$^+$) mice. The mean ± SEM are shown. Mann–Whitney test. Scale bars indicate 50 μm.

(L) NicheNet predicts potential ligands secreted by T/NK cell components, which regulate EC phenotype after DPAg treatment.

(M) Representative images of MECA79$^+$ cells that arise from either tdT$^-$ (LTβR wt, lightblue arrow) or tdT$^+$ (LTβR KO, yellow arrow) tumor vessels. White arrow indicates tdT$^+$ ECs. Scale bars indicate 20 μm.

Data are shown as mean ± SEM statistics were assessed by Mann-Whitney test (E, G, J) or two-way ANOVA (B, C, F, H, I at d14). The statistical analysis is referred to cDC2 (C). Data were pooled from two experiments (B, C, E-J).

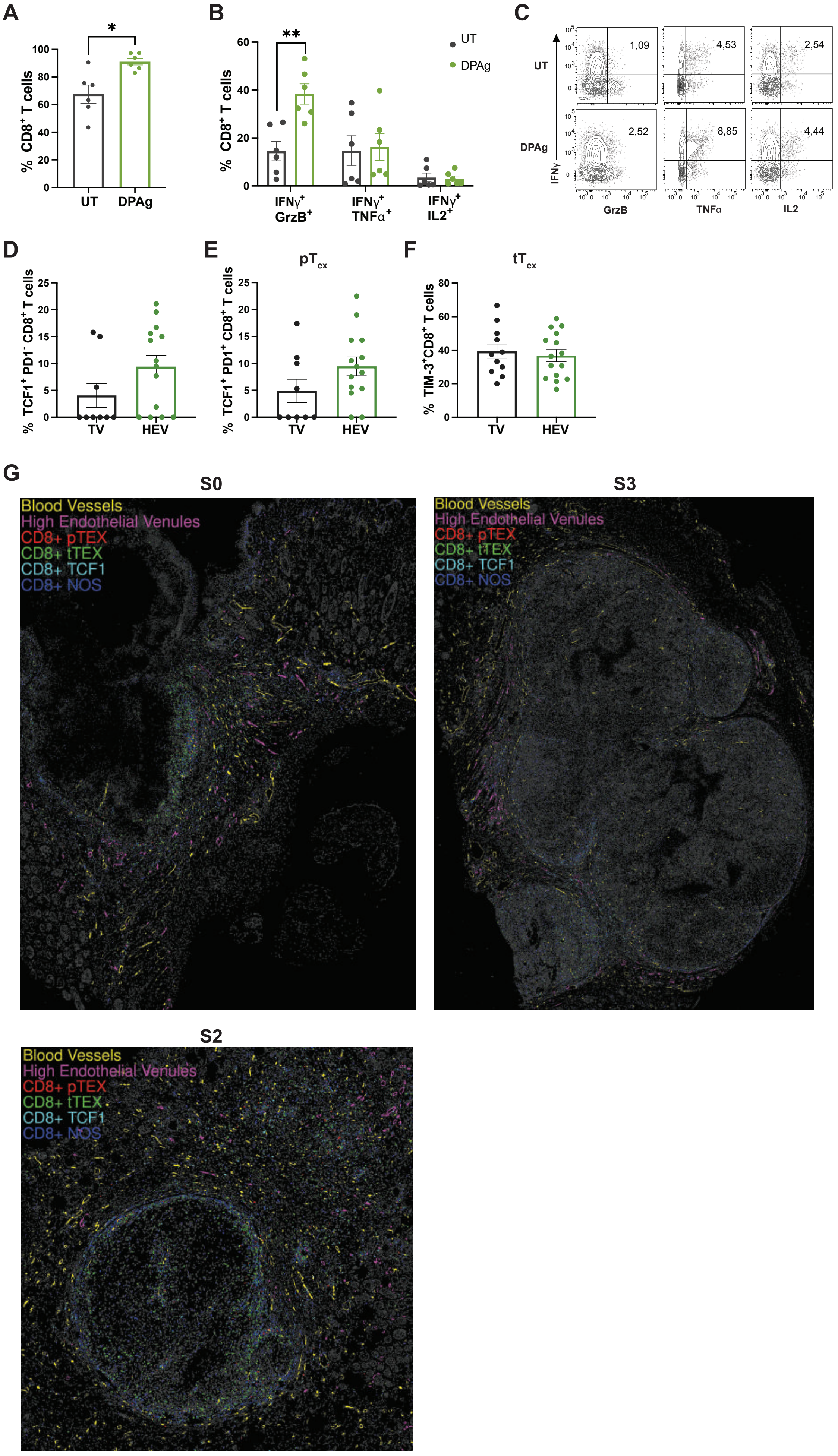
A
*
% CD8+ T cells
100
80
60
40
20
0
UT
DPAg
B
**
UT
DPAg
% CD8+ T cells
60
40
20
0
IFNγ+ GrzB+
IFNγ+ TNFα+
IFNγ+ IL2+
C
UT
DPAg
1,09
4,53
2,54
2,52
8,85
4,44
IFNγ
GrzB
TNFα
IL2
D
% TCF1+ PD1- CD8+ T cells
25
20
15
10
5
0
TV
HEV
E
pTex
% TCF1+ PD1+ CD8+ T cells
25
20
15
10
5
0
TV
HEV
F
tTex
% TIM-3+CD8+ T cells
80
60
40
20
0
TV
HEV
G
S0
S3
S2
Blood Vessels
High Endothelial Venules
CD8+ pTEX
CD8+ tTEX
CD8+ TCF1
CD8+ NOS

**Figure S6. TU-HEVs generate lymphocyte niches permissive for $PD1^{neg}$ and $PD1^+$ CD8 progenitor cells**

(A and B) Flow cytometry quantification of $PD1^+CD8^+$ T cells (A) and of $CD8^+$ T cells co-expressing IFNγ-GrzB or IFNγ-TNFα or IFNγ-IL2 (B) in UT (N = 6) and DPAg (N = 6) treated E0771 tumors. The mean ± SEM are shown.

(C) Representative flow cytometry dot plots of IFNγ-GrzB or IFNγ-TNFα or IFNγ-IL2 CD8 T cells of UT and DPAg-treated PyMT tumors.

(D-F) Quantification of $TCF1^+PD1^-CD8^+$ T cells (N = 9-14) (D), $pT_{EX}$ (N = 9-14) (E), and $tT_{EX}$ (N = 11-15) (F) 50 $\mu m^2$ around HEVs or other tumor vessels from E0771 frozen sections. The mean ± SEM are shown.

(G) Digital reconstruction of three MC38 tumor section stained with the MILAN multiplexing technique. $CD8^+$ $pT_{EX}$ (red) are $TCF1^+$ $PD1^+$ $TIM3^-$; $CD8^+$ $tT_{EX}$ (green) are $TCF1^-$ $PD1^+$ $TIM3^+$; $CD8^+$ TCF1 (light blue) are $TCF1^+$ $PD1^-$ $TIM3^-$. All the remaining $CD8^+$ T cells are identified as $CD8^+$ NOS (Not Otherwise Specified) (blue).

Mann-Whitney test (A, B, D-F). Data are pooled from two independent experiments (A, B, D-F).
